# Unveiling differential responses to UVB (305 nm) and UVC (275 nm) in cacao-infecting *Colletotrichum gloeosporioides* and *Pestalotiopsis* sp

**DOI:** 10.1101/2025.05.21.655292

**Authors:** Insuck Baek, Jae Hee Jang, Seunghyun Lim, Zhuangji Wang, Minhyeok Cha, Clint Magill, Moon S. Kim, Lyndel W. Meinhardt, Sunchung Park, Ezekiel Ahn

## Abstract

Sustainable control of microbial pathogens requires alternatives to chemicals, but physical methods like Ultraviolet-C (UVC) show variable efficacy linked to poorly understood pathogen-specific responses. Here, we investigate differential UVB/UVC responses in plant pathogenic fungi (*Colletotrichum gloeosporioides*, *Pestalotiopsis sp.*). Using hyperspectral imaging and machine learning, we dissect the physiological underpinnings of UV sensitivity. UVC proves more potent than UVB, with *Pestalotiopsis sp.* showing significantly higher resistance than *C. gloeosporioides* isolates. Crucially, hyperspectral signatures correlated with resistance, revealing UVC-induced photopigment changes, biochemical disruption, and oxidative stress markers in sensitive isolates, contrasting with minimal perturbation in the resistant isolate. Machine learning accurately decoded these complex phenotypes for classification. This understanding enabled enhanced inactivation via optimized pulsed UVC and synergistic sonication. We link distinct physiological states, non-invasively detected via hyperspectral imaging, to fungal UV resistance, demonstrating how integrating advanced phenotyping and machine learning provides a mechanistic basis for optimizing physical pathogen controls.

## Introduction

Fungal diseases pose a significant threat to global food security, causing substantial yield losses and economic damage in a wide range of crops. In the case of cacao (*Theobroma cacao* L.), fungal diseases such as frosty pod rot and black pod rot can devastate yields, leading to field abandonment and significant economic losses, which threaten the livelihoods of millions of smallholder farmers and, in turn, the global chocolate industry^1,2^. Furthermore, recent outbreaks of anthracnose caused by *Colletotrichum gloeosporioides* in Ghana have highlighted the vulnerability of cacao to emerging fungal threats^3^. Other foliar fungal pathogens of cacao include *Pestalotiopsis microspora* and *Nigrospora sphaerica*, causal agents of leaf infections and contributors to overall crop losses^4^. The *Pestalotiopsis* genus, although it is well known for its broad range of hosts and production of important secondary metabolites, remains largely unstudied and therefore lacks clarity, which can incur intragenus and intergenus confusion^5^.

Current control methods, primarily consisting of synthesized chemical fungicides, face growing challenges due to evolving resistance, environmental concerns, and consumer demand for safer food^6^. In this context, UV radiation, particularly in the UVC range (between 200 and 280 nm), has emerged as a promising alternative for controlling fungal pathogens due to its non-chemical nature, potential for targeted application, and ability to elicit plant defense mechanisms^7–9^. The most effective wavelength for UV disinfection is often considered to be 260 nm, which falls within the UVC range^10^; this wavelength is completely absorbed by the Earth’s atmosphere and does not naturally occur at the surface^11^. In practice, various studies suggest that the optimal wavelength can vary depending on the specific application, necessitating optimization for each case. For example, Janisiewicz et al. demonstrated that 222 nm far-UVC was three to ten times more effective than 254 nm UVC in inactivating certain strawberry fungal pathogens while leaving the host plant unharmed, stressing the importance of considering specific wavelengths and their interactions with target organisms^12^.

UVC radiation exerts its antifungal effects primarily by inducing DNA damage, forming photoproducts that disrupt cellular processes and can ultimately lead to cell death^13–15^. Through the absorption of UVC photons by DNA molecules, abnormal chemical bonds can form between adjacent thymine bases, forming thymine dimers^16,17^. These thymine dimers distort the DNA structure, which interferes with essential processes such as replication and transcription and ultimately hinders fungal growth and survival^18^. In addition to its direct antifungal activity, UVC radiation can also elicit plant defense responses by triggering signaling pathways that lead to the production of antimicrobial compounds against pathogen invasion^19–21^. This dual action of UVC radiation makes it a particularly attractive alternative to chemical fungicides for controlling fungal diseases in plants. UV-based technologies are being increasingly explored as a prospective form of fungal disease remediation in a variety of crops, including grapes^22,23^, strawberries^24^, and tomatoes^25,26^, with studies showing favorable results in reducing disease incidence and severity. As encouraging as the future for UV light treatment seems, the specifics have yet to be determined. Thus, further research is needed to determine optimal exposure times that disrupt fungal growth while minimizing any phytotoxicity that may occur in the plant host^21^.

The optimization of UV treatment necessitates the consideration of multiple factors such as the wavelength of UV light, exposure time, application methods, and an in-depth understanding of the complex interplay between UV radiation, fungal pathogens, and host plants. To address this need, this study employs a comprehensive approach that incorporates machine learning to analyze complex fungal responses to UV radiation and predict treatment efficacy. We investigated the effects of different UV wavelengths and exposure methods on the growth, morphology, and underlying physiological responses (via hyperspectral imaging) of two cacao-infecting fungi, *C. gloeosporioides* and *Pestalotiopsis* sp., with the aim of developing efficient and targeted UV-based control strategies. We hypothesized that UVC radiation, due to its higher energy photons, would be more capable of inhibiting the growth of cacao-associated fungal pathogens and inducing morphological changes than UVB radiation. Furthermore, we explored the capacity of pulsed UVC treatments to enhance the efficacy of UV-based control and investigated the synergistic effects of combining sonication with UVC exposure. This research has the potential to inform the development of multiple sustainable and productive UV-based disease management strategies for a wide range of crops, ultimately contributing to the ever-present issue of global food security.

## Results

### Phylogenetic analysis

Five fungal isolates (CGH5, CGH17, CGH34, CGH38, and CGH53), all initially labeled as *C. gloeosporioides*, were obtained from the Cocoa Research Institute of Ghana. These isolates were selected for this study to explore the diversity of fungal responses to UV radiation. To investigate the potential of UV-based control strategies for fungal pathogens in cacao and other crops, we first conducted in-house experiments using a custom-built UV box equipped with our custom-built UV monitor tools. To accurately identify the fungal isolates, we performed phylogenetic analysis using internal transcribed spacer (ITS) sequences (Fig. S1)^27^. The resulting phylogenetic tree revealed distinct clades within the *Colletotrichum* genus. CGH17, CGH34, CGH38, and CGH53 clustered with other *Colletotrichum* spp., including *C. gloeosporioides*. However, even within this group, some degree of divergence was observed. CGH34 grouped with *C. fructicola* and *C. chrysophilum*, while CGH38 and CGH53 were closely related to each other. Interestingly, CGH5 was placed in a distinct clade with *Pestalotiopsis* and *Neopestalotiopsis* spp., indicating that it does not belong to the *Colletotrichum* genus as initially presumed.

### Distinct growth patterns distinguish Colletotrichum gloeosporioides from Pestalotiopsis sp

Fungal morphology is a critical determinant of growth, development, and pathogenicity. This study investigated the morphology of four *C. gloeosporioides* isolates (CGH17, CGH34, CGH38, and CGH53) and one *Pestalotiopsis* sp. (CGH5). While CGH5, CGH17, and CGH53 were isolated from cacao, CGH34 and CGH38 were isolated from unidentified shade trees. Significant differences in colony morphology were observed between the isolates (Fig. 1a), indicating variations in growth patterns and potential ecological adaptations. Notably, CGH5 exhibited a significantly larger colony size (498.35 ± 9.24 mm^2^ mean ± SE) compared to the *C. gloeosporioides* isolates (CGH17 = 122.67 ± 4.1, CGH34 = 283.68 ± 6.35, CGH38 = 283.82 ± 11.38, and CGH53 = 223 ± 4.86). This observation is consistent with the distinct species classification of CGH5 as revealed by its ITS region amplification (Fig S3), which matched that of *Pestalotiopsis* sp. Within the *C. gloeosporioides* isolates, CGH17 exhibited significantly slower growth compared to the other isolates, as depicted in the representative images of CGH17 (Fig. 1b), CGH34 (Fig. 1c), and CGH5 (Fig. 1d) in Fig. 1. Colony shape-related traits (LWR; circularity; and IS & CG, presented in Fig. 1a, also revealed statistically significant differences, although some differences were slight. For instance, CGH17 exhibited the highest circularity (0.69 ± 0.0022), while CGH53 was considerably less circular (0.65 ± 0.003), and CGH34 fell in between (0.67 ± 0.0017).

**Fig. 1:**
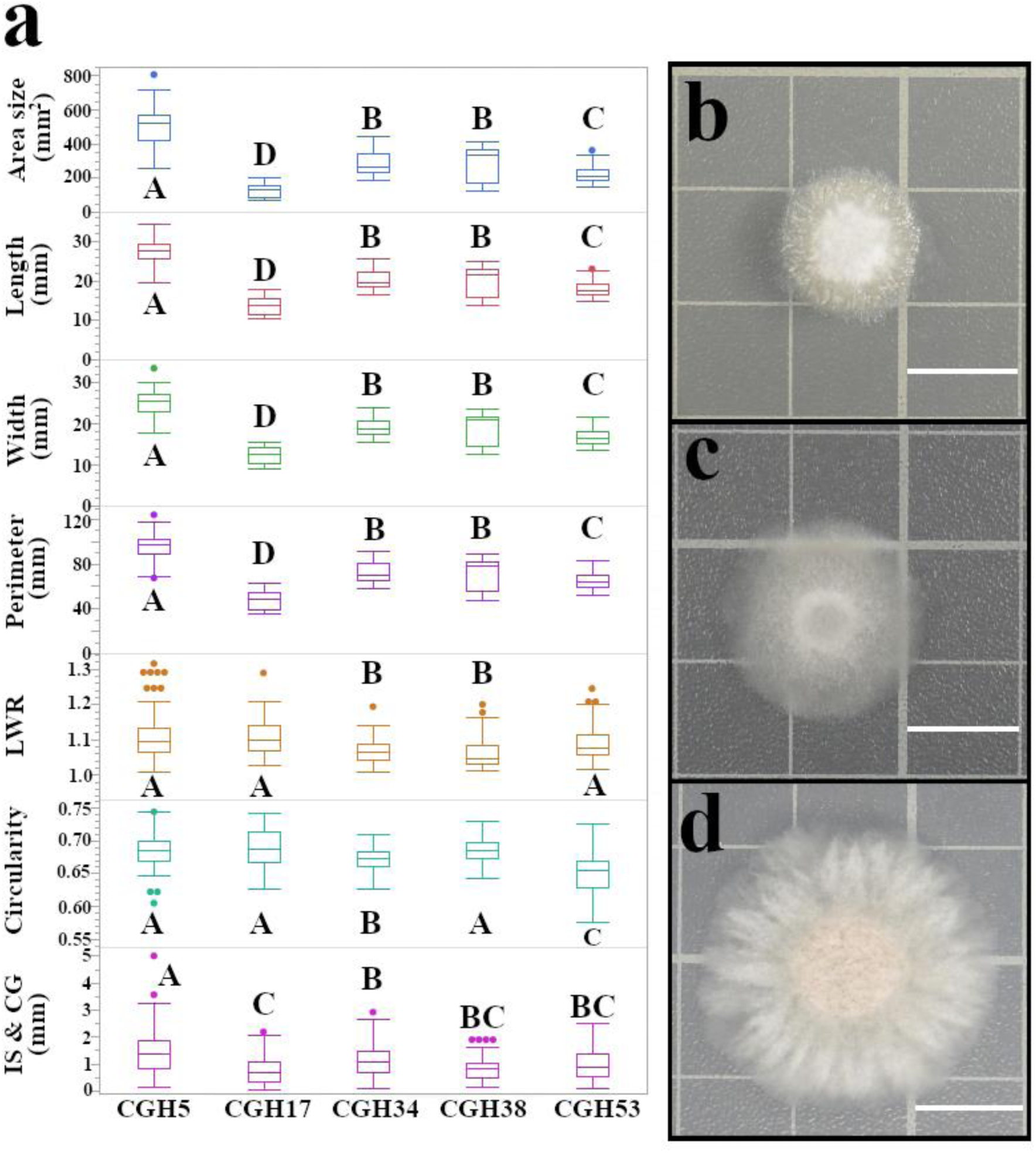
Distinct growth characteristics distinguish *C. gloeosporioides* isolates from *Pestalotiopsis* sp. **a** Quantitative analysis of colony morphology. *C. gloeosporioides* isolates (CGH17, CGH34, CGH38, and CGH53) and *Pestalotiopsis* sp. (CGH5) were cultured on potato dextrose agar (PDA) for 48 hours. Colony size and morphology were quantified and analyzed by one-way ANOVA with Tukey’s Honestly Significant Difference (HSD) post-hoc test (n = 457 colonies). Different letters indicate significant differences (*p* < 0.05). The majority of pairwise comparisons yielded *p* < 0.0001. **b-d** Representative colony morphology of CGH17 **b,** CGH34 **c,** and CGH5 **d**. Scale bar = 1 cm.

### UVB (305 nm) exposure reveals differential UVB sensitivity in fungi

As depicted in Fig. 2a, CGH5 exhibits a high level of resistance to UVB radiation (UVB intensity = 0.58 mW/cm^2^), evidenced by the minimal changes in colony size and morphology even after 30 minutes of exposure. Although a substantial initial decrease in colony size was observed when comparing the control (498.35 ± 9.24 mm^2^) (Fig 2b) to the colony size of 5-minute UVB-treated CGH5 (426 ± 4.86 mm^2^) (Fig 2c), further exposure did not lead to significant additional reductions at 10 minutes (414.76 ± 4.05 mm^2^) and 30 minutes (414.09 ± 4.93 mm^2^). All four *C. gloeosporioides* isolates presented varying degrees of sensitivity to UVB, with CGH38 displaying the most significant and gradual decrease in colony size (283.82 ± 11.38 mm^2^ in the control (Fig 2d) to 20.76 ± 4.86 mm^2^ after 30 minutes of UVB exposure (Fig 2e)) and CGH17 showing the smallest effect on colony size (122.76 ± 4.11 mm^2^ in the control to 90.33 ± 7.94 mm^2^ after 30 minutes of UVB exposure). Other size-related traits changed in an almost identical manner following UVB exposure; morphological traits such as circularity were altered in CGH17 (0.69 ± 0.0022 in the control to 0.56 ± 0.03 in the 30-minute UVB-treated) and CGH38 (0.68 ± 0.002 in the control to 0.29 ± 0.003 in the 30-minute UVB-treated), while other isolates maintained nearly the same level of circularity. The survival rate, or the presence of a visible colony at 48 hours of growth, showed that CGH5 remained at 1.0 following UVB 5, 10, and 30-minute treatments. Conversely, 30 minutes of UVB treatments caused CGH17 and CGH53 to drop down to 0.81 ± 0.089 and 0.83 ± 0.067, respectively, while CGH34 remained stable at 0.98 ± 0.015. Notably, CGH38 had the most significant reduction in survival rate, which plummeted to 0.40 ± 0.08.

**Fig. 2:**
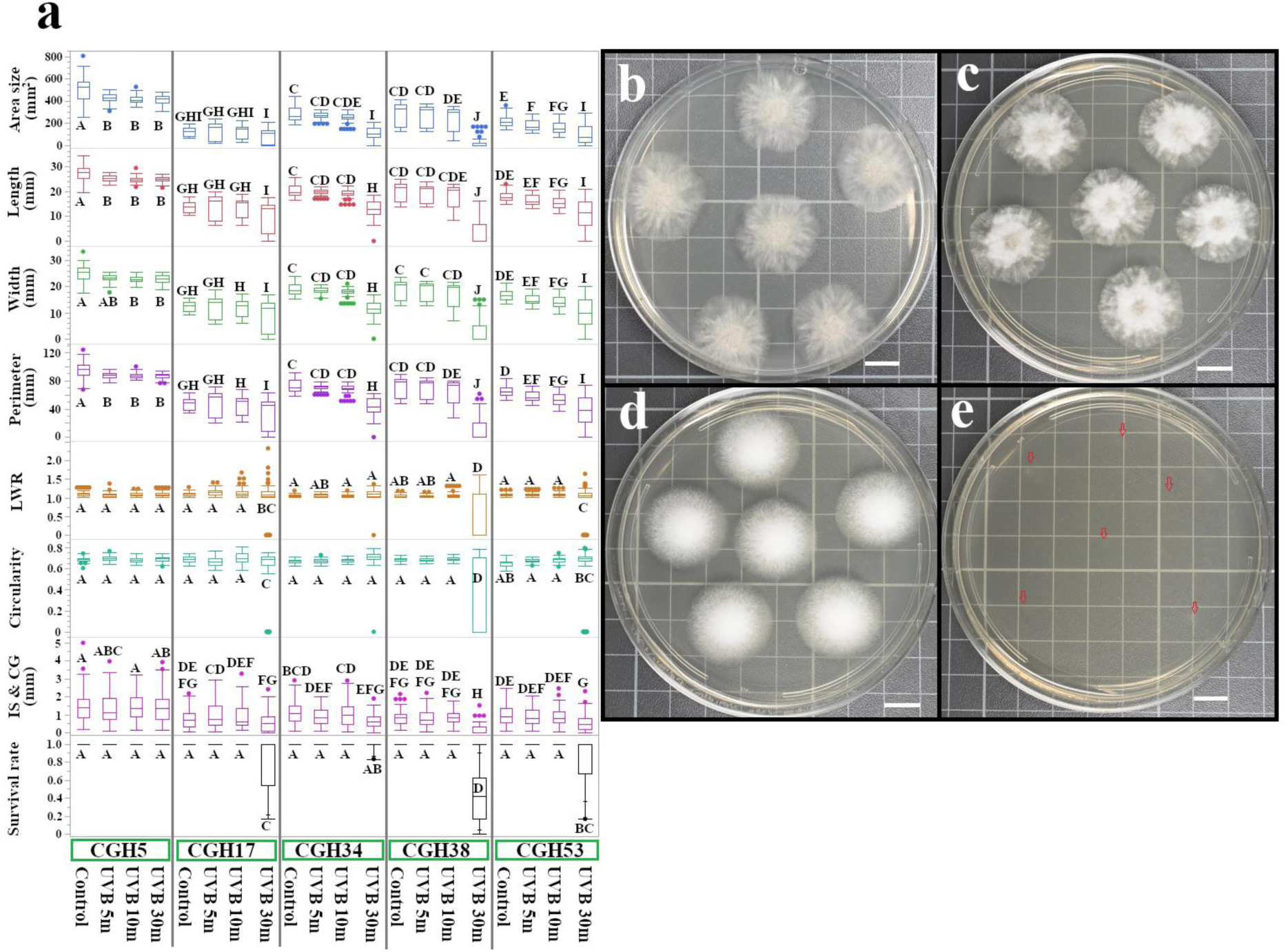
Differential UVB sensitivity of *C. gloeosporioides* and *Pestalotiopsis* sp. **a** Quantitative analysis of colony morphology following UVB exposure. *C. gloeosporioides* isolates (CGH17, CGH34, CGH38, and CGH53) and *Pestalotiopsis* sp. (CGH5) were exposed to UVB radiation for 5, 10, or 30 minutes. Colony size and morphology traits were measured and analyzed by one-way ANOVA followed by Tukey’s HSD post-hoc test (n = 1,515 colonies, including control). Different letters indicate significant differences (*p* < 0.05). The majority of pairwise comparisons yielded *p* < 0.0001. **b-e,** Representative images of *Pestalotiopsis* sp. (CGH5) and *C. gloeosporioides* (CGH38) colonies before (control) and after 30 minutes of UVB exposure. **b** CGH5 control; **c** CGH5, 30 min UVB; **d** CGH38 control; **e** CGH38, 30 min UVB. Red arrows indicate the conidia dispensed spots. Scale bar = 1 cm.

### UVC (275 nm) exposure inhibits fungal growth more effectively than UVB

Next, we examined the effects of UVC radiation on the fungal isolates (UVC intensity = 0.58 mW/cm^2^). The results indicated a significant decrease in colony size across all isolates, irrespective of their origin (Fig. 3a, CGH5 control displayed as a reference in Fig. 3b). UVC exposure was found to effectively inhibit the growth of *C. gloeosporioides* isolates within 5 minutes, as observed for CGH17 (Fig. 3c), CGH34 (Fig. 3d), CGH38 (Fig. 3e), and CGH53 (Fig. 3f). Noticeably, there was a substantial decrease in colony size after 30 minutes of UVC exposure (233.46 ± 11.31 mm^2^) compared to the control (498.35 ± 9.24 mm^2^), signifying a 53% reduction in colony size (Fig. 3b,g). This observation strongly aligns with the theory that 275 nm radiation causes substantial damage to the cacao-infecting fungal cells, leading to growth inhibition and cell death.

**Fig. 3:**
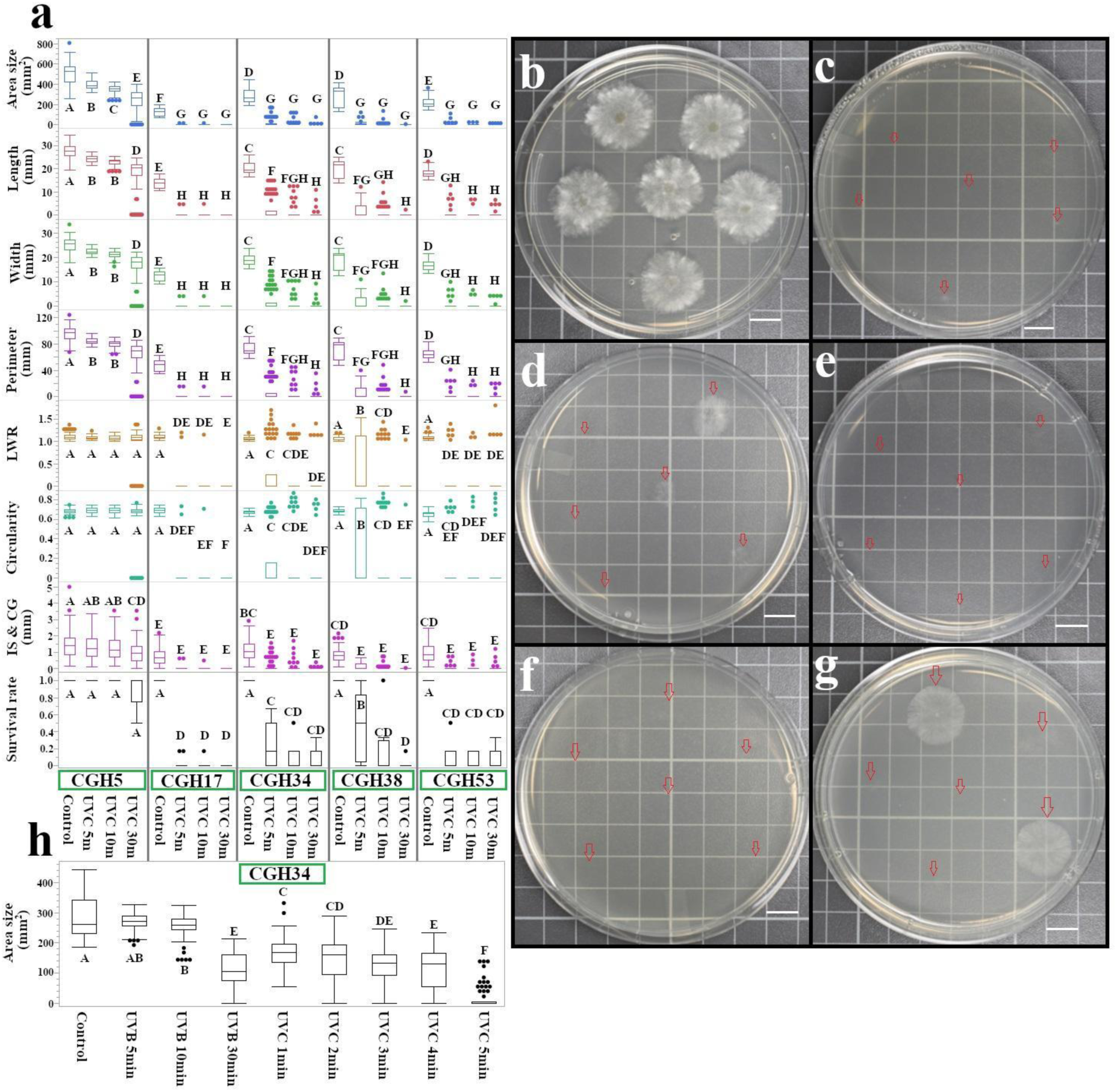
UVC exposure induces rapid and extensive morphological changes in cacao fungi pathogens. **a** Quantitative analysis of colony morphology following UVC exposure. *C. gloeosporioides* isolates (CGH17, CGH34, CGH38, and CGH53) and *Pestalotiopsis* sp. (CGH5) were exposed to UVC radiation for 5, 10, or 30 minutes (n = 1,531, including controls). Colony size and morphology were quantified and analyzed by one-way ANOVA with Tukey’s HSD post-hoc test. Different letters indicate significant differences (*p* < 0.05). The majority of pairwise comparisons yielded *p* < 0.0001. **b-g** Representative colony morphology before and after UVC exposure. Red arrows indicate conidia dispense spots. **b** CGH5 control; **c** CGH17, 5 min UVC; **d** CGH34, 5 min UVC; **e** CGH38, 5 min UVC; **f** CGH53, 5 min UVC; **g** CGH5, 30 min UVC. Scale bar = 1 cm. **h** Area size of *C. gloeosporioides* isolates following 1-4 minutes of UVC exposure. Control, UVB, and UVC 5 min treatments are included for comparison (n = 700, including other comparative groups).

Likewise, all other size-related traits, including the colony’s length, width, and perimeter, were significantly reduced, likely due to UVC-induced damage and growth inhibition. Shape-related traits such as LWR, circularity, and IS & CG also displayed significant differences upon exposure to UVC treatment.

Even with the exclusion of data points with no colony growth (area size 0) from the statistical analysis, significant differences in morphology across different fungal samples were still observed. For instance, CGH53 showed a change in circularity from 0.65 ± 0.003 in the control to 0.76 ± 0.042 after UVC exposure, while CGH5 maintained its shape over time. The survival rate of CGH5 remained relatively high at 0.86 ± 0.052, while it declined dramatically for all *Colletotrichum* isolates after 30 minutes of UVC-treatment; the survival rate was 0 for CGH17, 0.076 ± 0.03 for CGH34, 0.014 ± 0.014 for CGH38, and 0.076 ± 0.04 for CGH53.

To pinpoint the minimum length of UVC exposure time required to effectively kill *C. gloeosporioides* isolates, we conducted additional experiments with shorter exposure times (1-4 minutes) using CGH34. The results, as shown in Fig. 3h, indicate that UVC exposure gradually diminished the colony size from 283.68 ± 6.35 mm^2^ (control) to 168.79 ± 5.12 mm^2^ after 1 minute of UVC exposure, which further decreased to 145.24 ± 8.1 mm^2^ after 2 minutes, and then to 122.16 ± 6.69 mm^2^ and 113.1 ± 7.74 mm^2^ after 3 and 4 minutes of exposure, respectively. However, a dramatic reduction in colony size occurred at 5 minutes of UVC exposure, with a mean colony size of 17.37 ± 4.47 mm^2^. Notably, the resulting mean colony size after 4 minutes of UVC exposure nearly rivalled that of colonies subject to 30 minutes of UVB exposure (colony size of 114.14 ± 6.62 mm^2^); in this case, UVC was found to be more than seven times as effective as 300 nm UVB.

### Sonication enhances the efficacy of UVC treatment against *Pestalotiopsis* sp

We hypothesized that the relative resistance of CGH5 to UVC could be attributed to its robust cell wall structure, a trait that might provide physical protection against radiation damage. To investigate this, we employed sonication ((Elma Schmidbauer GmbH, Singen, Germany), 80 kHz, 100% power) to potentially disrupt the cell wall integrity of *Pestalotiopsis* sp. conidia prior to administration of UVC light.

Sonication alone had a limited influence on CGH5 growth, even with a prolonged treatment time of 30 minutes. While a reduction in colony size was observed (from 498.35 ± 9.24 mm^2^ in the control to 407 ± 6.3 mm^2^ after 1 minute of sonication and 215.98 ± 12.68 mm^2^ after 30 minutes), the survival rate remained largely unaffected (1.0 for 1-10 minutes of sonication and 0.85 for 30 minutes) (Fig. 4a). Interestingly, 30 minutes of sonication proved more effective in diminishing colony size than 30 minutes of UVB exposure, the latter of which maintained a survival rate of 1.0 and a colony size of 414.09 ± 4.93 mm^2^.

**Fig. 4:**
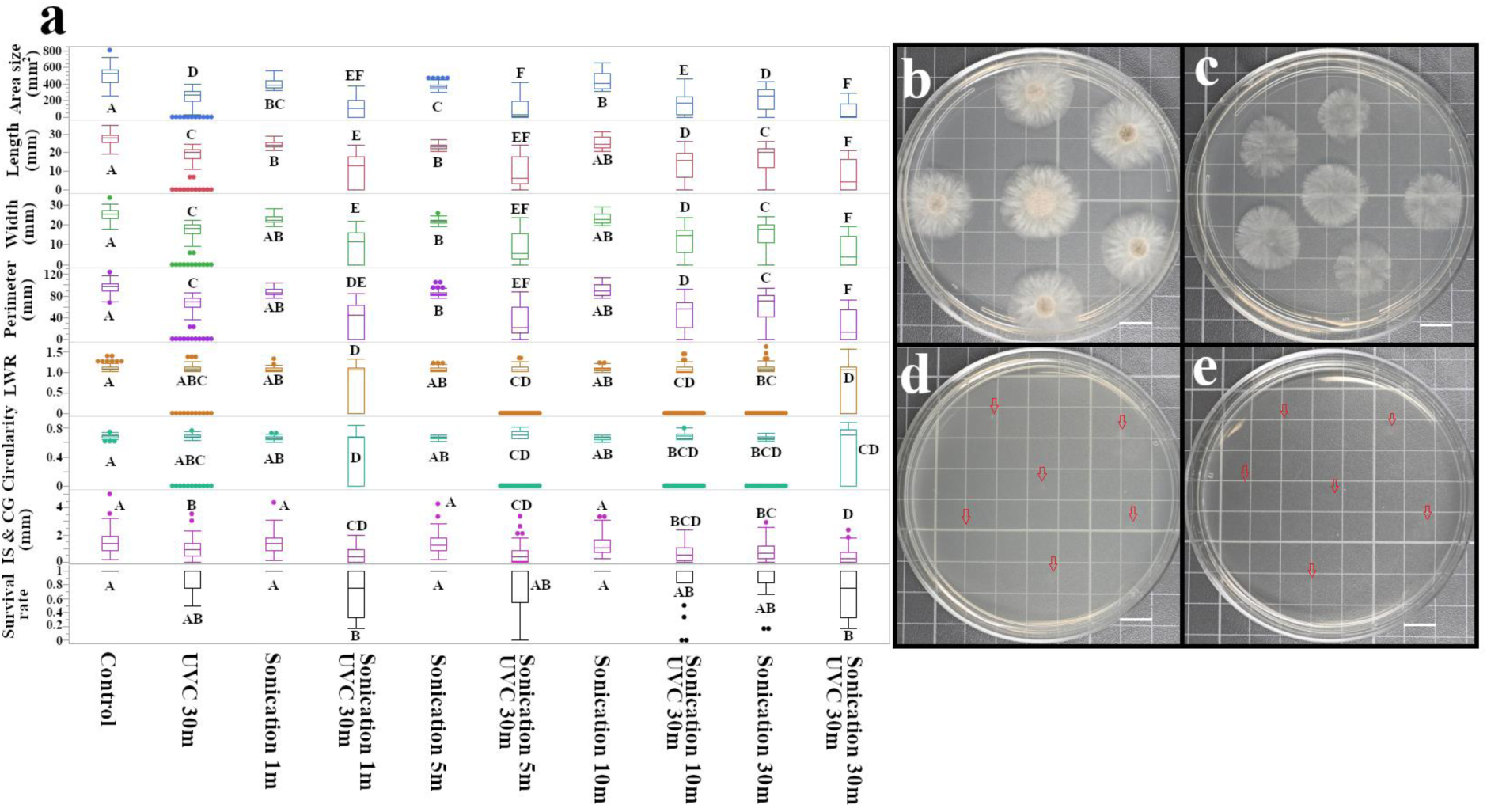
Synergistic effect of sonication and UVC exposure on the growth of *Pestalotiopsis* isolate. **a** Quantitative analysis of colony morphology following sonication and combined sonication/UVC treatment (n = 1,096 for morphological traits; n = 179 for survival rate). The *Pestalotiopsis* isolate (CGH5) conidia were sonicated for 1, 5, 10, or 30 minutes (80 kHz, 100% power) prior to inoculation on PDA. A subset of sonicated conidia was subsequently exposed to UVC radiation for 30 minutes. Colony size and morphology were quantified and analyzed by one-way ANOVA with Tukey’s HSD post-hoc test. Different letters indicate significant differences (*p* < 0.05) (n = 179). **b-e** Representative colony morphology of CGH5 following the indicated treatments. **b** Control (no treatment); **c** UVC 30 min; **d** Sonication 1 min + UVC 30 min; **e** Sonication 30 min + UVC 30 min. Scale bar = 1 cm.

While 30 minutes of UVC exposure alone significantly decreased colony size, it did not completely eliminate *Pestalotiopsis* sp., which maintained a survival rate of 0.89 (Fig. 4b,c). However, combining sonication and UVC had a synergistic effect, leading to a near-complete inhibition of *Pestalotiopsis* sp. growth (Fig. 4d). Remarkably, even 1 minute of sonication followed by 30 minutes of UVC exposure yielded a significant reduction in colony size (114.8 ± 11.4 mm^2^) and survival rate (0.68 ± 0.08). This synergistic effect was even more pronounced with longer sonication times (30 minutes) followed by UVC exposure, resulting in an area size of 73.63 ± 8.11 mm^2^ and a survival rate of 0.68 ± 0.06 (Fig. 4e).

### Pulsed UVC light exhibits isolate-specific effects on growth and morphology in cacao-infecting *Colletotrichum*

To investigate the effects of pulsed UVC light on fungal growth, we exposed *Colletotrichum* isolates CGH34 and CGH53 to 3-minute and 6-minute pulsed UVC treatments at varying frequencies (1 Hz, 10 Hz, and 20 Hz). The results of these treatments were compared to those of samples that underwent 3 continuous minutes of UVC exposure. As shown in Fig. 5a, the colony area of CGH34 under control conditions was 283.68 ± 6.35 mm^2^, which was brought down to 122.16 ± 6.69 mm^2^ following continuous UVC exposure. Interestingly, pulsed UVC at 1 Hz (201.08 ± 3.44 mm^2^) was less effective than continuous 3-minute UVC, while 10 Hz (51.79 ± 7.28 mm^2^) showed the most significant reduction, and 20 Hz (134.12 ± 4.97 mm^2^) fell in between (Fig.5 b-e). On the contrary, for CGH53, continuous UVC exposure (3 minutes) resulted in a near-complete growth inhibition (9.28 ± 1.12 mm^2^). Excluding 20 Hz, both 1 Hz and 10 Hz pulsed UVC treatments were less productive than continuous UVC in eliminating CGH53.

**Fig. 5:**
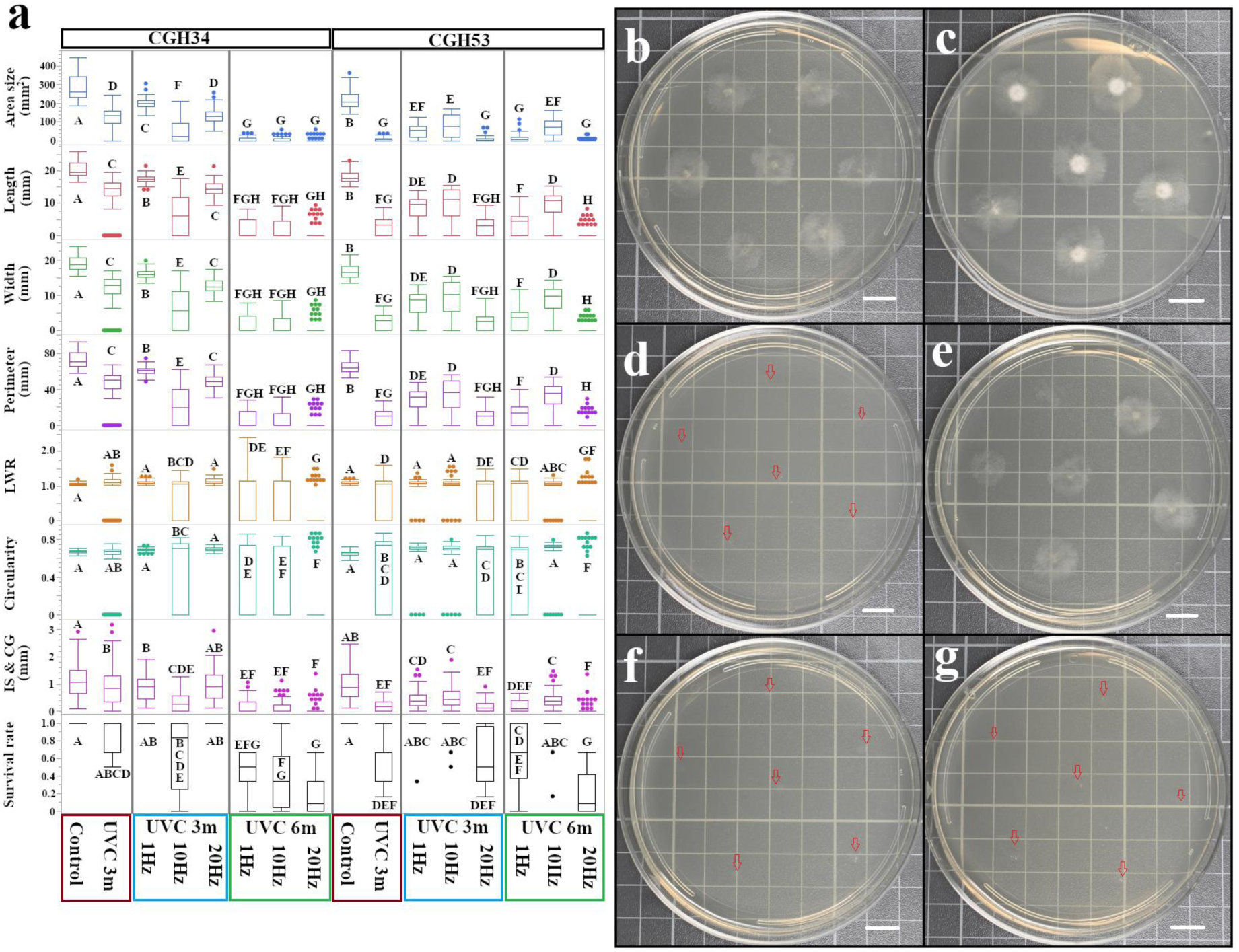
Pulsed UVC light enhances growth inhibition of cacao infecting *Colletotrichum* isolates. **a** Quantitative analysis of colony morphology following pulsed UVC exposure. *Colletotrichum* isolates (CGH34, CGH53) were exposed to UVC radiation for 3 minutes continuously or 3 minutes with varying pulse frequencies (1 Hz, 10 Hz, 20 Hz) (n = 1,229 for morphological traits; n = 204 for survival rate). To account for the reduced energy delivered during pulsed exposure, a 6-minute pulsed UVC treatment was also performed. Colony size and morphology were quantified and analyzed by one-way ANOVA with Tukey’s HSD post-hoc test. Different letters indicate significant differences (*p* < 0.05). **b-g** Representative colony morphology of CGH34 following the indicated UVC treatments. Red arrows indicate conidia dispense spots. **b** Control (3 min continuous UVC); **c** 3 min, 1 Hz; **d** 3 min, 10 Hz; **e** 3 min, 20 Hz; **f** 6 min, 10 Hz; **g** 6 min, 20 Hz. Scale bar = 1 cm.

To account for the reduced energy delivered during pulsed UVC exposure, we also tested 6-minute pulsed treatments. In CGH34, 6-minute pulsed UVC led to further shrinking of colony size compared to the 3-minute continuous treatment, with 20 Hz showing the greatest effect (4.2 ± 1.29 mm^2^) (Fig. 5f,g). Similarly, in CGH53, 6-minute pulsed UVC at 20 Hz resulted in the smallest colony size (2.59 ± 0.78 mm^2^), while the 6-minute 10 Hz treatment was surprisingly less potent than even the 3-minute continuous UVC. These results indicate that pulsed UVC can be more effective than continuous UVC in inhibiting fungal growth, but the optimal pulse frequency and duration may vary depending on the fungal isolate. This isolate-specific response could be attributed to differences in DNA repair mechanisms, pigmentation, cell wall composition, or other cellular characteristics that influence the fungi’s sensitivity to UVC radiation. Pulsed UVC also impacted colony morphology. For instance, differences in circularity were observed between CGH34 exposed to continuous UVC (0.58 ± 0.0017) and 1 Hz pulsed UVC (0.69 ± 0.002). These findings suggest that pulsed UVC not only affects fungal growth but also disrupts fungal development and morphogenesis.

The survival rates of the isolates also varied with UVC treatment. For CGH34, 3 minutes of continuous UVC resulted in a survival rate of 0.86 ± 0.051, while 3 minutes of both 1 Hz and 20 Hz pulsed UVC were of no consequence (1.0). However, doubling the length of the pulsed UVC treatments to 6 minutes led to a frequency-dependent reduction in survival, with 20 Hz being the most effective (0.17 ± 0.062). CGH53 showed a similar trend, with 20 Hz pulsed UVC most successfully lessening the survival rate.

### Multivariate and t-SNE analyses reveal distinct responses to UV treatments and phylogenetic influence on morphological traits

To explore the relationships between UV treatment, fungal isolate, and fungal colony morphology, we conducted a multivariate analysis encompassing principal component analysis (PCA), Pearson correlation, and hierarchical clustering. The biplot (Fig. 6a) displays the distribution of fungal isolates subjected to various UV treatments along the first two principal components (PCs), which account for 53.7% of the total variation. Notably, UVB treatments cluster closely together, as do UVC treatments. Furthermore, UVC treatments of 1, 2, and 4 minutes are closely grouped, suggesting a similar effect on fungal morphology. Control samples (no UV treatment) cluster near 5-minute UVB and 10-minute UVB treatments, pointing to a minimal impact on fungal growth within this timeframe. The 30-minute UVC treatment group closely aligns with 10-minute UVC, 1-minute sonication with 30-minute UVC, and 5-minute sonication with 30-minute UVC treatments, hinting at a potential threshold for UVC’s effect on fungal morphology.

**Fig. 6:**
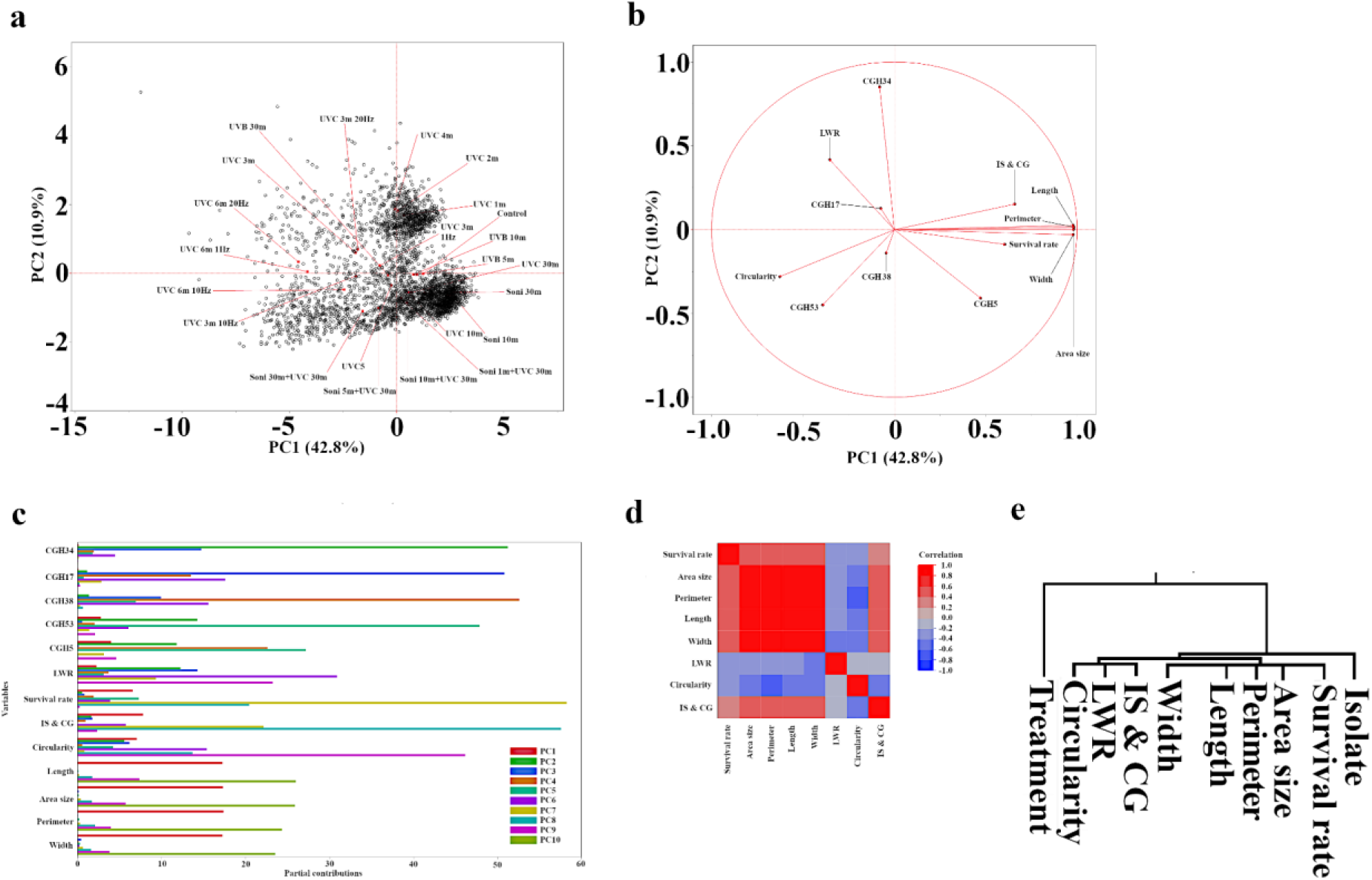
Principal component analysis and correlation of morphological traits reveal distinct responses to UV treatments. **a** Biplot of principal component (PC) scores for fungal isolates subjected to various UV treatments. Data points represent individual colonies, and treatments are labeled to illustrate separation along PC1 and PC2. **b** Loading plot of PC1 and PC2, showing the contribution of morphological traits and fungal isolates to the observed variation. **c** Partial contribution plot of PC1-PC10, illustrating the contribution of fungal isolates and morphological traits to each principal component. Note the similarity among size-related traits and the distinct contribution of survival rate. **d** Pearson correlation matrix of morphological traits. All pairwise comparisons were statistically significant (*p* < 0.0002). Analysis was performed on log-transformed data to address non-normality. **e** Hierarchical clustering analysis of UV treatment, isolate, and morphological traits using Ward’s linkage method.

The loading plot (Fig. 6b) depicts the contribution of morphological traits and fungal isolates to the observed variation. Unsurprisingly, size-related traits (area, perimeter, length, and width) positively correlate with and contribute similarly to PC1. Conversely, shape-related traits (IS & CG, LWR, and circularity) demonstrate dissimilarity, indicating differential responses to UV treatments. Survival rate aligns closely with size-related traits, implying a compelling link between growth inhibition and overall fungal survival. All five isolates show varied responses, although CGH53 and CGH38 express similar trends, mirroring their phylogenetic proximity. Fig. 6c presents the partial contribution plot of PC1-PC10, further showcasing the contribution of fungal isolates and morphological traits to each principal component. Size-related traits depict consistent patterns across PCs, while circularity, IS & CG, and LWR exhibit some similarities, particularly with strong peaks in PC6. LWR and circularity are also influenced by PC9. Survival rate shows distinct peaks in PC7 and PC8, similar to IS & CG, but with an inverse relationship in PC7 and PC9, as is reflected in Fig. 6b. Apart from CGH5 and CGH53, two isolates that share a strong contribution from PC5, the tested isolates display diverse responses, with dominant PCs differing greatly.

The Pearson correlation matrix of morphological traits (Fig. 6d) reveals that all pairwise comparisons are statistically significant (*p* < 0.0002). Notably, survival rate conveys strong positive correlations with area-related traits (0.54 to area size, perimeter, length, and width), while negatively correlating with LWR (−0.26) and circularity (−0.35). IS & CG shows a slight positive correlation (0.29) with the survival rate. Hierarchical clustering analysis (Fig. 6e) using Ward’s linkage method reveals distinct groupings. UV treatments form an outgroup, shape-related traits cluster together, size-related traits group with survival rate, and isolates form a separate cluster.

Furthermore, we investigated the relationship between the evolutionary history of the isolates and their morphological traits. Patristic distances among the five isolates, calculated using the Neighbor-Joining method based on their ITS sequences, were correlated with morphological traits. We found significant positive correlations between patristic distance and colony area (0.35, *p* = 0.0064), perimeter (0.35, *p* = 0.0073), length (0.36, *p* = 0.0055), width (0.36, *p* = 0.0059), and IS & CG interaction (0.38, *p* = 0.0028). A moderate positive correlation was also observed between patristic distance and LWR (0.28, *p* = 0.031). These results suggest that more closely related isolates tend to exhibit more similar morphological traits, underlining the influence of evolutionary history on fungal morphology and potentially its response to UV stress.

Fig. 7 presents a t-SNE visualization exploring the morphological responses of *C. gloeosporioides* isolates and a *Pestalotiopsis* isolate to different UV treatments. Fig. 7a displays the overall grouping of isolates across all treatments, with green dots representing the CGH5 isolate and black dots representing other *C. gloeosporioides* isolates. Shifting focus to the effects of UVC radiation, Fig. 7b highlights isolates exposed to 30 minutes of UVC (cyan dots). A noticeable shift in clustering compared to the overall grouping in Fig 7a indicates that UVC 30 minute exposure induces significant morphological/survival rate changes compared to the other treatments. Furthermore, the scattered distribution of cyan dots suggests variability in UVC sensitivity among the isolates. Fig. 7c compares the morphological responses of CGH5 isolates with (red dots) and without (green dots) sonication treatments followed by 30 minutes of UVC exposure. The overall separation between clusters, albeit with some overlap, suggests that sonication moderately impacts UVC-induced morphological changes. Fig. 7d provides a comprehensive view by examining the overall fungal response to all treatments, including UVB 5-, 10-, 30-minute (violet dots), UVC 5-, 10-, 30-minute (cyan dots), and control (yellow dots) conditions.

**Fig. 7:**
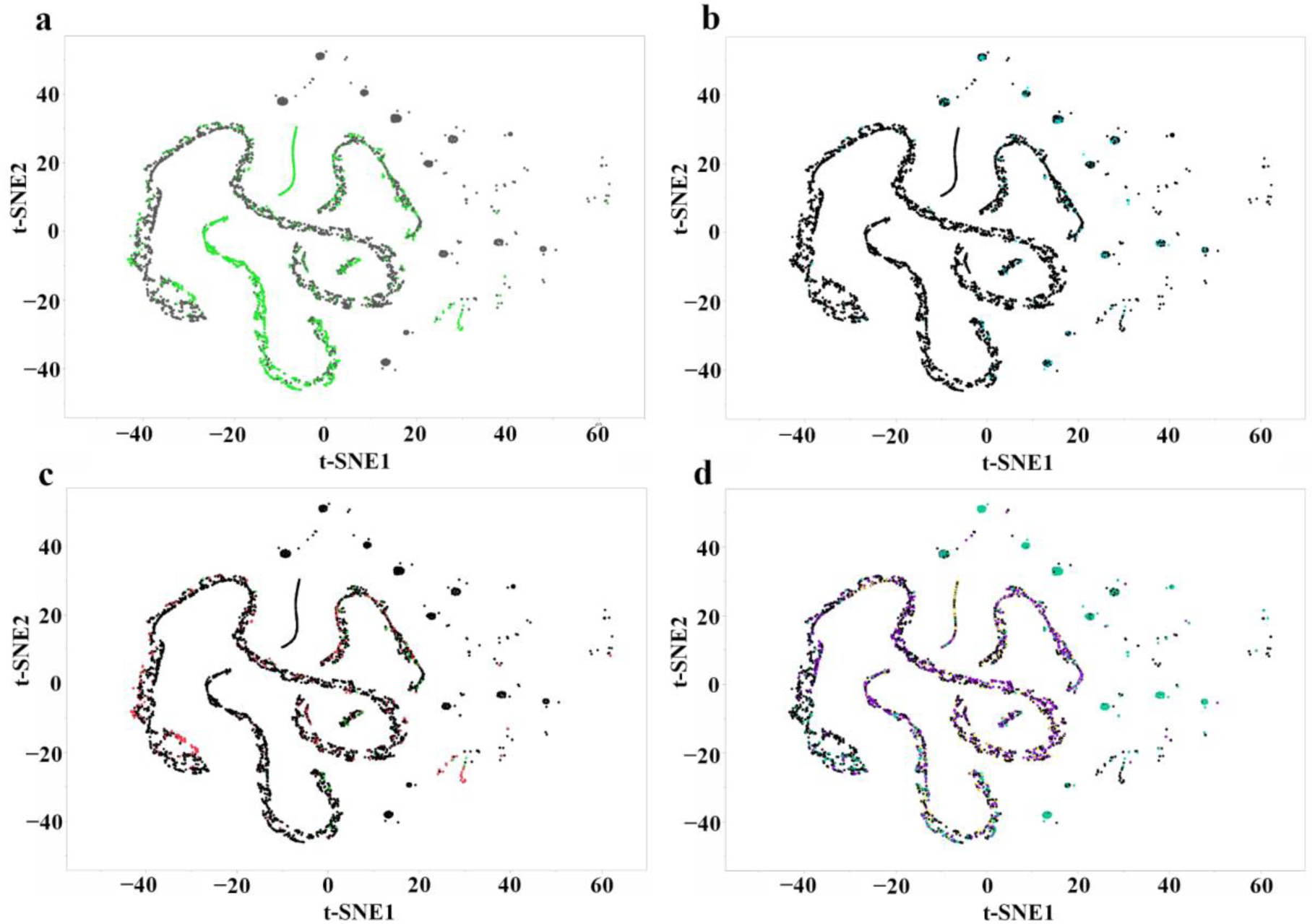
t-Distributed stochastic neighbor embedding visualization of *C. gloeosporioides* isolates and a *Pestalotiopsis* isolate based on morphological traits following exposure to different stress treatments. **a** Overall clustering pattern of CGH5 (green) and other *C. gloeosporioides* isolates (black) across all treatments. **b** Highlighting the morphological response of isolates to 30 minutes of UVC exposure (cyan). **c** Comparison of CGH5 isolates with sonication for at least 1 minute (red) versus no sonication (green) after 30 minutes of UVC exposure. **d** Overall response to all treatments, including UVB 5, 10, 30 min (violet), UVC 5, 10, 30 min (cyan), and control (yellow) conditions for all isolates.

### Hyperspectral Imaging Reveals Differential Fungal Responses to UVC Stress

Visible and near-infrared (VIS-NIR) reflectance spectra (430-1000 nm) were compared before (baseline) and 24 hours after 30 minutes of UVC exposure for isolates CGH53 and CGH5 (Fig. 8a,b). For the UVC-sensitive isolate CGH53, a notable decrease in reflectance, corresponding to increased absorption, was observed across the measured spectral range after UVC treatment compared to its baseline spectrum (Fig. 8a). In contrast, the UVC-resistant isolate CGH5 exhibited minimal changes in its VIS-NIR reflectance spectrum following UVC exposure (Fig. 8b).

**Fig. 8.**
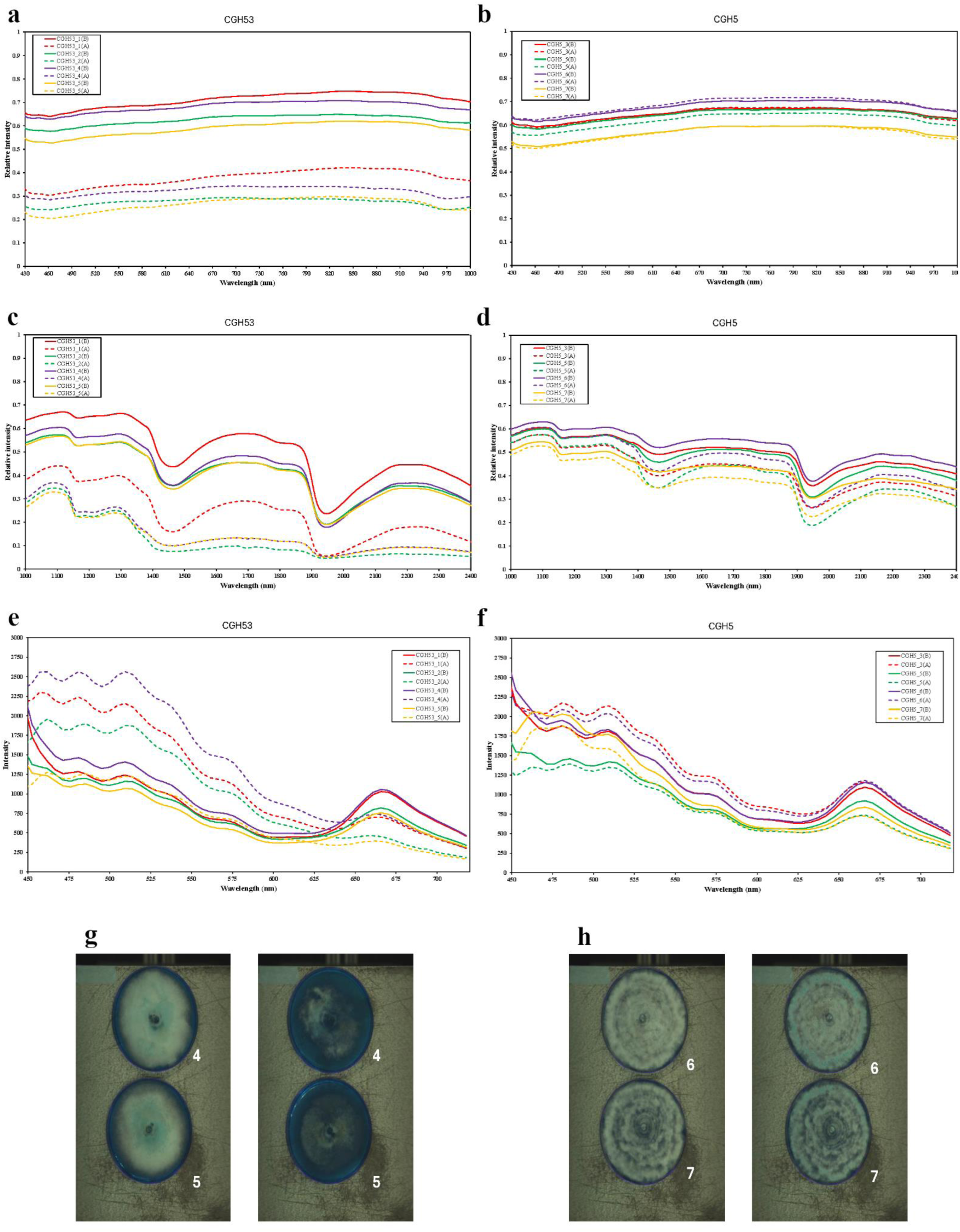
Differential hyperspectral responses and SWIR imaging of *C. gloeosporioides* (CGH53) and *Pestalotiopsis* sp. (CGH5) to UVC radiation. Spectral profiles and representative short-wave infrared (SWIR) reflectance images of fungal isolates CGH53 (left column, **a, c, e, g**) and CGH5 (right column, **b, d, f, h**) before (baseline) and 24 hours after 30 minutes of UVC exposure (275 nm, 0.58 mW/cm²). **a-f** Mean spectral profiles from four biological replicates (n=4); solid lines represent baseline spectra, dashed lines represent spectra 24h after UVC exposure. **a, b** Visible and Near-Infrared (VIS-NIR) reflectance spectra (430-1000 nm), showing decreased reflectance in CGH53 **a** but minimal change in CGH5 **b** post-UVC. **c, d** SWIR reflectance spectra (1000-2400 nm), indicating notable changes in water (∼1450, 1900 nm) and lipid (∼1700-1750 nm) associated bands for CGH53 **c** but stability in CGH5 **d**. **e, f** Fluorescence emission spectra (450-720 nm, 365 nm excitation), revealing significantly elevated emission in CGH53 **e** versus negligible change in CGH5 **f** after UVC treatment. **g, h** Representative SWIR reflectance images comparing the appearance of fungal colonies before UVC (left images) and 24h after UVC (right images) for **g** CGH53 (sample ID 4 and 5 shown) and **h** CGH5 (sample ID 6 and 7 shown), illustrating visual changes, particularly the darkening observed in CGH53 post-treatment.

Short-wave infrared (SWIR) reflectance spectra (1000–2400 nm) were analyzed pre- and post-UVC treatment for CGH53 and CGH5 to assess biochemical changes (Fig. 8c,d). In CGH53, distinct spectral changes were observed after UVC exposure, notably increased absorption (lower reflectance) in regions associated with water content (around 1450 nm and 1900 nm) and variations in absorption features within the 1700–1750 nm range, associated with C-H bonds in lipids/organic structures (Fig. 8c). Conversely, isolate CGH5 displayed minimal alterations in its SWIR reflectance spectrum after UVC treatment, with absorption levels at the water and lipid-associated bands remaining largely unchanged compared to its baseline (Fig. 8d).

Fluorescence emission spectra (450-720 nm, excited by 365 nm UV-A) were also compared before and after UVC treatment (Fig. 8e,f). Following UVC exposure, CGH53 exhibited significantly elevated fluorescence intensity across the 500–600 nm range compared to its baseline emission (Fig. 8e). Heightened emission was particularly apparent around 530-560 nm and also observed around 500 nm. These spectral regions correspond to known emission ranges for various endogenous fluorophores, including flavins, melanin, carotenoids, and lipofuscin-like compounds (Matveeva et al., 2024). In stark contrast, CGH5 showed only negligible changes in fluorescence emission intensity and spectral shape after UVC treatment compared to its baseline (Fig. 8f).

Representative images derived from the SWIR reflectance data visually supported these findings; CGH53 showed distinct visual changes, including darkening of the colonies 24 hours post-UVC treatment compared to baseline (Fig. 8g), whereas CGH5 colonies appeared largely unchanged between the two time points (Fig. 8h). Raw hyperspectral values for each wavelength (VIS-NIR, SWIR, and fluorescence) are available in Supplementary Data 2, and images illustrating the appearance of the culture plates before and after treatment are provided in Supplementary Data 3.

### Machine learning accurately predicts fungal isolate classification based on UV treatment and morphological traits

To further investigate the relationship between UV treatment, fungal isolates, and morphological traits, we employed machine learning techniques to predict fungal isolate classification. Eight different models were evaluated (Table S1): Bootstrap Forest^28^, Decision Tree^29^, Nominal Logistic^30^, Naive Bayes^31^, Generalized Regression Lasso^32^, K-Nearest Neighbors (k-NN)^33^, Neural Boosted^34^, and Support Vector Machines (SVM)-Radial Basis Function (RBF) model^35^.

Using a stratified random sampling approach, each isolate was represented by 509 samples, which were then split into training and validation sets split into an 80:20 ratio. Bootstrap Forest and Neural Boosted achieved over 70% accuracy for training and validation sets. Specifically, Bootstrap Forest achieved 83.04% accuracy in the training set and 71.6% in the validation set, and Neural Boosted achieved 73.49% and 73.2% accuracy, respectively.

A detailed analysis of the Neural Boost model (NTanH(3) NBoost(20)) revealed that CGH5 had the highest classification accuracy (0.97 in the validation set), while CGH34 (54.9%) and CGH53 (56.3%) displayed comparatively lower accuracy (Table S2). This variation in accuracy may reflect differences in the sensitivity and morphological responses of different isolates to UV treatments. It may also stem from CGH5 potentially belonging to a completely different fungal genus, while CGH34 and CGH53 are phylogenetically closer and therefore more similar and harder to distinguish.

The magnitude of different features in the Neural Boost model was also assessed (Table S3). UV treatment conditions were the most dominant feature, followed by area size, length, and width. IS & CG, circularity, and survival rate held less weight in the classification model. The high importance of UV treatment condition (total effect of 0.705) aligns with its significant influence on fungal morphology and survival. Visual representations of the Receiver Operating Characteristic (ROC) and Area Under the Curves (AUC) for the different models are available in Fig. S2a and b. Fig. S2c and d display the ROC and AUC curves for different isolates in the Neural Boost model.

### Predicting fungal colony area size using machine learning

To further explore the relationship between UV treatment, fungal isolate, and colony morphology, we employed machine learning models (Boosted Tree^36^, Bootstrap Forest^28^, Decision Tree^29^, Least Squares^37^, Stepwise^38^, Generalized Regression Lasso^32^, k-NN^33^, Neural Boosted^34^, and SVM-RBF model^35^) to predict fungal colony area size based solely on UV treatment conditions and fungal isolate information. This analysis excluded direct size and shape-related features as predictors, focusing on the predictive power of UV treatment and isolate type alone. Even with the exclusion of direct size and shape measurements, several models, including Boosted Tree, Neural Boosted, and Decision Tree, achieved high predictive accuracy (R-squared > 0.84) in both training and validation sets (Table S4 and Fig. S3).

## Discussion

This study investigated the effects of different UV wavelengths and exposure methods on the growth and morphology of cacao-associated fungi, aiming to explore the potential of UV-based approaches for controlling fungal pathogens in agriculture. Our findings reveal a complex interplay between UV treatment, fungal isolate/strain, and morphological responses, accentuating the importance of tailoring UV-based strategies to specific pathogens and their susceptibilities. Notably, we observed significant differences in the sensitivity of *C. gloeosporioides* and *Pestalotiopsis* sp. to UVB (305 nm) and UVC (275 nm) radiation. The initial reduction in colony size of CGH5 followed by a plateau in growth inhibition suggests that the isolate may possess a highly efficient DNA repair mechanism, protective secondary metabolites^39^, a fast growth habit that allows it to outpace the detrimental effects of UVB, or an adaptive response to UVB stress^40^, allowing it to mitigate the damage stemming from prolonged UVB exposure. The contrasting responses of each isolate highlight the need to consider both the fungal species and the specific UV wavelength when developing control strategies. The phylogenetic analysis emphasizes the diversity of fungal isolates associated with cacao and the importance of accurate identification for understanding their biology and developing effective control strategies. The inclusion of CGH5, despite its unexpected placement within the *Pestalotiopsis* clade, provided valuable insights by introducing significant variety to the responses of the fungal species to UV radiation.

Our observation that UVC (275 nm) was more effective in inhibiting fungal growth aligns with prior findings by Wan et al., who reported significant membrane damage in fungal spores of *Aspergillus niger*, *Penicillium polonicum*, and *Trichoderma harzianum* upon exposure to UV-LEDs emitting at 280 nm and a 265/280 nm combination^41^. This damage was linked to increased intracellular reactive oxygen species (ROS) levels, ultimately contributing to enhanced inactivation of the fungal spores^41,42^. These findings informed our choice of UVC wavelength (275 nm), as we sought to investigate the outcome of administering UV radiation (near the 280 nm range) on plant pathogenic fungi to forge future agricultural applications.

Interestingly, previous studies have also revealed that dark conditions did not promote significant DNA repair in the fungal spores they studied, implying that the damage induced by UV exposure may persist in the absence of light^41,43^. This observation is consistent with our findings that the effects of UV exposure on fungal morphology were still evident after 48 hours of incubation in the dark. This lack of dark repair could be advantageous for UV-based control strategies, as it indicates that a single exposure could inflict lasting inhibitory influences on fungal growth.

The observation of changes in fungal morphology runs parallel to findings from a recent work demonstrating the ability of UV mutagenesis to intensify the antagonistic activity of *Trichoderma* spp. against plant fungal pathogens^44^. In the referenced study, UV-induced mutations in *T. virens* and *T. asperellum* led to improved biocontrol capabilities, signifying that UV exposure can both directly inhibit fungal growth and induce changes that affect the fungi’s interactions with other organisms. This finding raises the intriguing possibility that pulsed UVC treatment could potentially enhance the efficacy of biocontrol agents like *Trichoderma* spp., either by directly increasing their growth or by inducing mutations that amplify their antagonistic activity.

The synergistic effect observed between sonication and UVC exposure in inhibiting *Pestalotiopsis* sp. growth (Fig. 4) warrants further discussion. While sonication alone had a limited impact on fungal survival, its combination with UVC resulted in a significant reduction in both colony size and survival rate. Perhaps sonication improves the efficacy of UVC treatment by disrupting the fungal cell wall, thereby increasing the penetration of UVC radiation and causing greater cellular damage. This would have serious implications for developing combined treatment strategies against UVC-resistant fungal pathogens, particularly in cases where increasing UVC intensity alone might be detrimental to the host plant. The capacity of sonication to supplement the antifungal effects of other treatments is supported by Campaniello et al., who demonstrated that combining sonication with heat (thermo-sonication) adequately inactivated fungal spores of *Penicillium* spp. and *Mucor* spp. in distilled water^45^. They found that the power of the sonication treatment primarily influenced the antifungal activity of thermo-sonication. Similarly, sonication pretreatment has also been shown to significantly enhance the degradation of grape pomace by fungal enzymes, suggesting that sonication can increase the accessibility of biomass components to enzymatic attack^46^. From a practical perspective, considering the probable limitations of prolonged sonication and UVC exposure in field applications, the observation that even 1 minute of sonication can synergistically boost UVC efficacy is highly encouraging. This finding suggests that a brief sonication pretreatment could significantly scale down the required UVC exposure time, theoretically minimizing any adverse effects on the host plant while maximizing the efficacy of fungal pathogen control. Importantly, our preliminary observations indicate that cacao leaves can tolerate UVC exposure for 10 minutes at the intensity used in this study without incurring significant damage (no data shown). Thus, UVC treatments could potentially be applied directly to cacao plants without causing deleterious consequences for the host, although further research is needed to confirm this. Additional investigation is needed to optimize the sonication parameters, devise the exact method of sonication, apply said method to the appropriate host plant growth stage, and explore the concerted effects it may have with other antifungal agents, namely UVC irradiation, for controlling cacao-associated fungi.

Our results demonstrate that pulsed UVC treatments, particularly at 10 Hz and 20 Hz, led to significantly greater reductions in colony size and survival rates compared to continuous UVC exposure, even when the total energy delivered was comparable. This enhanced efficacy could be attributed to the disruption of fungal DNA repair mechanisms. The intermittent “off” periods in pulsed UVC exposure might prevent the fungus from sufficiently repairing the damage caused during the “on” periods, leading to a greater accumulation of DNA damage and ultimately cell death. This hypothesis is supported by previous studies demonstrating the ability of pulsed UV light to inactivate various microorganisms, including plant fungal pathogens ^47^. Scott et al. demonstrated that pulsed UVC treatment effectively controlled *Botrytis cinerea* in tomatoes, likely due to the activation of multiple defense pathways and the suppression of genes involved in fruit softening^48^. A subsequent study showed that pulsed UVC significantly reduced *B. cinerea* in lettuce, with the added benefit of shorter treatment times compared to continuous UVC^49^.

The visible reflectance spectra provided the initial view of differential responses to UVC stress. For CGH53, comparing spectra before and after UV exposure established a significant change post-treatment, perhaps due to the fungus producing more UV-absorbing pigments (i.e., melanin) or suffering surface structural changes after irradiation (Fig. 8a). Fungal melanin is known to absorb across UV-visible wavelengths and is often upregulated as a protective response to UV damage^50,51^. The increased absorption (lower reflectance) in CGH53 after UV is consistent with melanization, which would help shield fungal cells from further UV harm. This finding is supported by prior observations that fungi often accumulate melanin in cell walls under UV stress as a defense ^51^. Structural damage (like spore or cell surface changes) could also contribute to lower reflectance by scattering less light. In contrast to these changes, CGH5’s visible reflectance remained relatively stable after the same UV treatment (Fig. 8b). The before-and-after spectra for CGH5 overlap closely, with no significant drop in the blue/UV region, indicating that CGH5 neither produces appreciable additional pigment nor expresses symptoms of surface damage from the UV exposure. The stability of CGH5’s VIS spectrum suggests that this fungus either already possesses protective pigmentation or structural features, or it effectively resisted the UV-induced changes that affected CGH53. Some fungi have constitutive levels of melanin or other photoprotective compounds present even without induction^50^. It is possible that CGH5 had such pre-existing defenses (for example, a naturally higher melanin content or perhaps a thicker cell wall, a hypothesis consistent with its increased resistance being overcome by sonication combined with UVC) that buffered it against UV stress.

Further probing the fungi’s biochemical integrity, SWIR spectroscopy provided insights into changes related to water content, lipid composition, and structural integrity. This study analyzed the SWIR reflectance spectra of CGH53 and CGH5 before and after UV exposure to assess their biochemical responses (Figs. 8c and 8d). For CGH53, notable spectral changes were observed post-UV treatment, with a significant decrease of reflectance (increased absorption) in both water-associated absorption bands around 1450 nm and 1900 nm and lipid-associated bands between 1700–1750 nm. The increased absorption at 1450 nm and 1900 nm may stem from moisture loss or redistribution within fungal cells^52^. The 1700–1750 nm absorption features, associated with lipid structures, displayed variation, implying possible lipid peroxidation or alterations in cell membrane composition due to UV exposure^52,53^. In contrast, CGH5 displayed minimal spectral changes in SWIR reflectance before and after UV exposure, indicating that it maintained stable water content and lipid integrity (Figs. 8g and 8f). The absence of significant shifts in water and lipid bands indicates that CGH5 was more resistant to UV-induced dehydration and oxidative damage. This stability may be attributed to intrinsic protective mechanisms such as efficient membrane repair, pre-existing pigmentation, or antioxidant defenses.

Fluorescence spectroscopy further illuminated distinct oxidative stress responses. UV-induced stress in sensitive CGH53 significantly elevated emission (approx. 500–600 nm) relative to baseline, corresponding to known fluorophores including flavins, protective pigments (melanin, flavonoids), and oxidative damage markers like lipofuscin (Fig. 8e)^54^. This indicates the accumulation of both protective compounds and metabolites resulting from cellular oxidative damage. In stark contrast, resistant CGH5 demonstrated minimal fluorescence changes, suggesting negligible induction of these stress markers (Fig. 8f). Taken together, the multi-modal hyperspectral analyses consistently depict the sensitive CGH53 undergoing significant UV-induced physiological stress (pigment changes, biochemical disruption, oxidative metabolite accumulation), whereas the resistant CGH5 maintains remarkable stability, indicative of robust pre-existing or rapidly deployed protective mechanisms.

The successful application of machine learning underscores its potential for interpreting complex biological responses and informing control strategies. High predictive performance (Bootstrap Forest, Neural Boosted) in classifying isolates based on UV-induced morphology demonstrates potential for rapid diagnostic tools, leveraging the models’ ability to capture non-linear phenotypic relationships. Furthermore, accurately predicting colony area size using only isolate identity and UV treatment confirms these factors as key determinants of fungal growth responses, aiding efficacy assessments for optimized UV applications. While this study links phenotypic and physiological responses to UV resistance, precisely pinpointing the contributions of specific mechanisms (e.g., DNA repair efficiency, baseline pigmentation, metabolic shifts, signaling pathway interactions) requires further investigation. Future research integrating molecular techniques (e.g., transcriptomics, proteomics) with our hyperspectral and machine learning framework promises a deeper understanding of these complex interactions and the development of highly targeted control strategies.

In summary, this work demonstrates that integrating multi-spectral hyperspectral imaging with machine learning provides a powerful, non-invasive approach to dissect the physiological underpinnings of differential UV radiation resistance in fungi. By linking distinct spectral signatures, reflecting pigmentation, biochemical integrity, and oxidative stress, to pathogen susceptibility, we establish a mechanistic basis for observed variations in UVC efficacy. This understanding moves beyond empirical screening, enabling the rational design of optimized physical control strategies, such as pulsed UVC delivery and synergistic treatments involving sonication, tailored to overcome specific resistance mechanisms. Looking forward, this integrated ‘phenomics-to-prediction’ paradigm holds significant promise for broader applications in microbiology, from studying diverse microbial stress responses to engineering targeted, sustainable control methods for agricultural, environmental, or even clinical challenges, reducing reliance on conventional chemical agents. Validating the specific molecular pathways suggested by these physiological fingerprints via genomic and transcriptomic analyses represents a key next step in fully leveraging this approach.

## Methods

### UV box design and construction

A schematic of the UV treatment system designed for controlled exposure of fungal samples is shown in Fig. S4. The system incorporates UVB, UVC, and combined UVB+UVC LED modules mounted on the top frame, along with a LiDAR camera for capturing 3D depth data. A programmable linear stage moves the sample through the exposure area based on predefined settings, while the LiDAR simultaneously records structural features such as volume and shape. LED intensity is adjustable via a digital dimmer switch. The entire process is automated through in-house software, with integrated safety sensors and visual indicators to ensure operator protection.

### Fungal isolates and DNA extraction

Five fungal isolates, initially identified as *Colletotrichum gloeosporioides* (CGH17, CGH34, CGH38, and CGH53), were obtained from the Cocoa Research Institute of Ghana. An additional isolate, CGH5, was initially included in this group but later identified as *Pestalotiopsis* sp. All isolates were maintained on potato dextrose agar (PDA) at 24°C in the dark. For DNA extraction, isolates were cultured on 90 mm PDA plates for 10 days. Mycelium was harvested using a sterile inoculation loop and transferred to a microcentrifuge tube. Genomic DNA was extracted using the Zymo Quick-DNA Fungal/Bacterial Microprep kit according to the manufacturer’s instructions.

### PCR amplification of ITS region

The ITS region of the ribosomal DNA was amplified using ITS1 and ITS4 primers^27^. PCR reactions were performed in a 50 μl volume containing 25 μl of GoTaq G2 Green Master Mix, 2X (Promega), 2 μl of each primer (10 μM), 5 μl of template DNA, and 18 μl of nuclease-free water. The PCR cycling conditions were as follows: initial denaturation at 95°C for 2 minutes, followed by 35 cycles of denaturation at 95°C for 1 minute, annealing at 48°C for 30 seconds, and extension at 72°C for 45 seconds. A final extension step was performed at 72°C for 7 minutes. A Veriti™ Thermal Cycler (Applied Biosystems, Waltham, Massachusetts, USA) was used for PCR amplification. PCR products were verified using gel electrophoresis, which was performed in 2% agarose 1x TAE gel containing GelRed 10,000x in DMSO (Thomas Scientific, Swedesboro, New Jersey, USA). The verified amplicon PCR products were then sent to Psomagen (Rockville, Maryland, USA) for Sanger sequencing.

### Phylogenetic analysis

The ITS sequences of the fungal isolates were analyzed to determine their phylogenetic relationships. In order to generate consensus sequences for each isolate, *de novo* assemblies were performed in the Geneious Prime software using the acquired ITS1 and ITS2 sequences from Psomagen. Regions of poor sequencing accuracy were manually removed. The resulting sequences were compared to those in the NCBI GenBank database using BLASTn to identify closely related sequences (sequences of the 5.8S rRNA gene, flanked by the ITS1 and ITS2 regions in fungi). The ITS sequences and reference sequences were aligned using MUSCLE^55^ with Super 5 in Geneious Prime. The aligned sequences were then exported to MEGA software ^56^ for phylogenetic analysis. A phylogenetic tree was constructed using the Neighbor-Joining method combined with an interior-branch test and 1,000 bootstrap replications. The tree was condensed to exclude bootstrap values less than 80% and then transferred to FigTree for further clarity- and aesthetic-related editing.

### Fungal inoculum preparation for UV treatment

Five fungal isolates, initially identified as *C. gloeosporioides* (CGH17, CGH34, CGH38, and CGH53) and one isolate identified as *Pestalotiopsis* sp. (CGH5), were obtained from the Cocoa Research Institute of Ghana. Isolates were maintained on potato dextrose agar (PDA) at 24°C under dark conditions. To prepare fungal inoculum, isolates were cultured on PDA plates for 10 days. Conidia were harvested by flooding the plate with 10 ml of sterile distilled water containing 0.05% Tween 20 and gently scraping the surface of the colony with a sterile glass spreader. The spore suspension was filtered through four layers of sterile cheesecloth to remove mycelial fragments. The concentration of spores in the suspension was determined using a hemocytometer and adjusted to 1 × 10^6^ conidia/ml using sterile distilled water. A sterile 10 μl pipette was used to dispense the fungal inoculum onto fresh PDA plates. For each isolate, 6-7 spots of 10 μl inoculum were placed onto the PDA plate.

### UV treatment

Following inoculation, PDA plates were immediately placed into a custom-built UV box equipped with either UVC (275 nm) or UVB (305 nm)-LED lights. The intensity of both UVB and UVC light sources was monitored using a radiometer RMD (Opsytec Dr. Gröbel GmbH, Ettlingen, Germany) and adjusted to 0.58 mW/cm^2^. Fungal isolates were exposed to UVB radiation for 5, 10, or 30 minutes. For UVC treatments, isolates were exposed for 5, 10, or 30 minutes, with an additional set of experiments using shorter exposure times (1, 2, 3, and 4 minutes) for *C. gloeosporioides* isolate CGH34. Control plates were not exposed to UV radiation. Following UV treatment, plates were incubated at 24°C in the dark for 48 hours to allow for colony growth.

### Sequential sonication and UVC exposure

To investigate the combined effects of sonication and UVC radiation, *Pestalotiopsis* sp. isolate CGH5 was subjected to sonication treatment prior to UVC exposure. A sonication bath (Elma Schmidbauer GmbH, Singen, Germany) operating at 80 kHz and 100% power was used for sonication treatments. Fungal spores were suspended in 1 ml of sterile distilled water in a 1.5 ml microcentrifuge tube and placed in the sonication bath for 1, 5, 10, or 30 minutes. Following sonication, 10 μl of the spore suspension was dispensed onto fresh PDA plates as described above. A subset of the sonicated spores was then exposed to UVC radiation for 30 minutes as described in the UV Treatment section. Control plates were not exposed to either sonication or UVC radiation. All plates were incubated at 24°C in the dark for 48 hours to allow for colony growth.

### Pulsed UVC treatment

*C. gloeosporioides* isolates CGH34 and CGH53 were subjected to pulsed UVC treatments. Pulsed UVC light was generated using a UVC-LED lamp (275 nm) with a built-in pulse function. Isolates were exposed to 3-minute and 6-minute pulsed UVC treatments with varying frequencies (1 Hz, 10 Hz, and 20 Hz). These treatments were compared to a continuous 3-minute UVC exposure. Following UV treatment, plates were incubated at 24°C for 48 hours in the dark to allow for colony growth. Fungal colonies were then imaged using a Nikon Z8 camera with a Z 20mm lens (Nikon, Tokyo, Japan) and a Kaiser R2N CP stand (Kaiser Fototechnik, North White Plains, NY) to ensure consistent image capture. Morphological measurements were taken from the captured images.

### Image analysis

Fungal colony morphology was characterized using SmartGrain software (version 1.3) ^57^. Although primarily designed for seed morphology analysis, SmartGrain was adapted for this study due to its ability to accurately quantify key fungal colony morphological traits, including area, length, width, length-weight relationship (LWR), perimeter, circularity, and the distance between the intersection of length and width (IS) and center of gravity (CG) (IS & CG). Individual colonies were manually identified and measured in each image using SmartGrain. A total of 4,769 colonies from 796 plates were analyzed. Each experiment was replicated at least three times to ensure reproducibility (Raw data available in Supplementary Data 1).

### Statistical Analysis

Statistical analyses were performed using JMP Pro 17 software (SAS Institute Inc.)^58^. One-way analysis of variance (ANOVA) was used to compare means between different treatment groups. Tukey’s Honestly Significant Difference (HSD) post-hoc test was used to identify significant pairwise differences between means. Pearson correlation analysis was used to assess correlations between morphological traits. Hierarchical clustering analysis was performed using Ward’s linkage method. Principal component analysis (PCA) was conducted to explore the relationships between UV treatments, fungal isolates, and morphological traits. All PCA analyses were performed on log-transformed data to address non-normality in some traits. Statistical significance was defined as *p* < 0.05. t-SNE analysis was performed in JMP Pro 17 using all quantitative morphological traits as input variables. Algorithm parameters: output dimensions = 2, perplexity = 30, maximum iterations = 1,000, initial principal component dimensions = 50, convergence criterion = 1e−8, initial scale = 0.0001, Eta (learning rate) = 200, inflate iterations = 250, and random seed = 123.

To assess the relationship between the five isolates’ phylogeny and morphology, we performed additional Pearson’s correlation analysis between the patristic distances and morphological traits. For this analysis, categorical variables, such as UV treatment conditions, were converted to one-hot encoding, and time of UV exposure was treated as a numerical variable. All morphological traits, including one-hot encoded variables, were log-transformed to improve normality and homoscedasticity. Spearman’s rank correlation analysis was then performed to assess the relationships between the log-transformed patristic distances and the log-transformed morphological traits. Statistical significance was defined as *p* < 0.05.

### Hyperspectral imaging of fungal response to UVC stress

Based on observed differential resistance to UV treatment, isolates *Pestalotiopsis sp.* CGH5 (resistant) and *C. gloeosporioides* CGH53 (sensitive) were selected for hyperspectral analysis to investigate underlying physiological responses. Fungal inocula were prepared and dispensed onto PDA plates as previously described. Four biological replicates, consisting of independent plates for each selected isolate, were prepared. Each plate was first imaged using the hyperspectral systems described below to acquire baseline (pre-treatment control) spectral data. Immediately following this initial imaging, the same plates were exposed to UVC radiation (275 nm) for 30 minutes at an intensity of 0.58 mW/cm² using the custom UV treatment system. After UVC exposure, the plates were incubated at 24°C in the dark for 24 hours. Following this incubation period, the plates were imaged a second time with the hyperspectral systems under identical conditions to acquire post-treatment spectral data. This paired approach allowed for direct comparison of spectral changes induced by UVC treatment within each replicate plate.

Three types of hyperspectral images, including fluorescence, visible and near-infrared (VNIR) reflectance, and short-wave infrared (SWIR) reflectance, were obtained using custom-developed imaging systems (Fig. S5). A VNIR hyperspectral line-scan imaging system was employed to acquire both reflectance and fluorescence images. The system includes a camera unit, a programmable motorized linear stage (Velmex, Inc., Bloomfield, NY, USA) for sample movement, and an illumination unit designed to accommodate two illumination sources specific to each imaging mode. Reflectance imaging utilized illumination from a 150-W DC quartz-tungsten halogen lamp (Dolan Jenner, Boxborough, MA, USA), coupled via optical fibers to generate two thin line lights directed at the instantaneous field of view (IFOV). Fluorescence imaging employed UV excitation from a pair of custom UV-A units (365 nm), each containing four 10-W LEDs (LedEngin, San Jose, CA, USA) and integrated fans for cooling. The camera unit consists of a 14-bit electron-multiplying charge-coupled device (EMCCD) camera (Luca DL 604M, Andor Technology, South Windsor, CT, USA), combined with a VNIR hyperspectral imaging spectrograph (400–1000 nm, VNIR Hyperspec, Headwall Photonics, Fitchburg, MA, USA), a lens with a 23-mm focal length, and a 60-μm slit. Using this system, reflectance hyperspectral images were collected in the spectral range of 430–1000 nm, and fluorescence images were acquired within the 450-720 nm range. Additional technical information on this VNIR imaging system is available in Kim et al. (2011).

The SWIR hyperspectral line-scan imaging system operates similarly to the VNIR system but differs primarily in its camera and illumination components. The instantaneous field of view (IFOV) is illuminated by two custom-designed illumination units, each containing four 150-W gold-coated halogen lamps equipped with MR16 reflectors, extruded heat sinks, and cooling fans. The camera unit comprises a 16-bit mercury cadmium telluride (MCT) array detector paired with a 25-mm focal length lens and an imaging spectrograph (Hyperspec-SWIR, Headwall Photonics, Fitchburg, MA, USA). Using this setup, SWIR reflectance images were captured in the spectral range of 1000–2400 nm^59^.

### Machine learning for fungal isolate classification

To investigate the relationship between UV treatment, fungal isolates, and morphological traits, machine learning techniques were employed to predict fungal isolate classification. A dataset was compiled, consisting of morphological measurements (area, perimeter, length, width, LWR, circularity, and IS & CG) and survival rates for each fungal isolate under different UV treatments. Eight different machine learning models were evaluated: Bootstrap Forest, Decision Tree, Nominal Logistic Regression, Naive Bayes, Generalized Regression Lasso, k-NN, Neural Boosted, and SVM-RBF. Data preprocessing involved a stratified random sampling approach to ensure equal representation of each isolate (509 samples per isolate, totaling 2,545 samples). The dataset was then split into 80% training and 20% validation sets. JMP Pro 17 software was used for model training and evaluation. Model performance was assessed based on classification accuracy in both the training and validation sets (random seed = 1). The importance of different features in the best-performing model (Neural Boosted) with the settings NTanH(3) NBoost(20) was also evaluated. ROC curves and AUC values were generated to visualize model performance.

### Machine learning for predicting fungal colony area size

To further explore the relationship between UV treatment, fungal isolate, and colony morphology, machine learning models were employed to predict fungal colony area size. Unlike the previous analysis, this analysis focused solely on the predictive power of UV treatment conditions and fungal isolate information, excluding direct size and shape measurements as predictors. Nine different machine learning models (Boosted Tree, Bootstrap Forest, Decision Tree, Least Squares Regression, Stepwise Regression, Generalized Regression Lasso, k-NN, Neural Boosted, and SVM-RBF) were evaluated for their ability to predict colony area size. The dataset used for this analysis included the fungal isolate and the UV treatment condition as input features, and the colony area size as the output variable. The dataset was split into 80% training and 20% validation sets. JMP Pro 17 software was used for model training and evaluation. Model performance was assessed based on the coefficient of determination (R-squared) in both the training and validation sets with the model screening function (random seed = 1). The importance of different features in the best-performing model (Neural Boosted) with the settings NTanH(3) NBoost(20) was also evaluated. ROC curves and AUC values were generated to visualize model performance.

### Statistical information

Statistical analyses were performed using JMP Pro 17. All experiments were independently replicated at least three times (biological replicates), with multiple colonies often measured per replicate (technical replicates). Key analyses included one-way ANOVA with Tukey’s HSD post-hoc tests, Pearson and Spearman correlation, PCA on log-transformed data, Hierarchical Clustering (Ward’s method), and t-SNE. Significance was defined as *p* < 0.05. Exact sample sizes (n), specific test statistics, and exact *p*-values for all analyses are reported within the Results section, figure legends, or supplementary materials. Data visualizations and any error bars shown are defined within the corresponding figure legends. Statistics related to machine learning models are detailed separately in the Methods section.

## Supporting information

Supplemental Figures and Tables, and will be used for the link to the file on the preprint site.

## Acknowledgements

We are also grateful to the reviewers for their constructive feedback. This work is supported by the U.S. Department of Agriculture, Agricultural Research Service, In-House Projects No. 8042-21220-258-000-D and 8042-21000-303-000-D. Mention of any trade names or commercial products in this article is solely for the purpose of providing specific information and does not imply recommendation or endorsement by the U. S. Department of Agriculture. USDA is an equal opportunity provider and employer, and all agency services are available without discrimination.

## Author contribution

E.A. designed the experiments. E.A. directed the overall study. L.W.M., M.S.K., and E.A. obtained funding for its execution; I.B., E.A., and S.L. performed machine learning analysis. S.L. and E.A. conducted bioinformatics and statistics. I.B., Z.W., C.M., M.C., S.P. provided resources. S.P., M.S.K., L.W.M., and E.A. provided supervision. J.H.J., C.M., and M.C. provided support developing methodology. All authors contributed to the writing and editing of this manuscript.

## Data availability

Phenotypic data are available in the Supplementary Data 1. Raw hyperspectral data (VIS-NIR, SWIR, Fluorescence) are provided in Supplementary Data 2. Representative images of culture plates are available in Supplementary Data 3. Further data are available from the corresponding author upon reasonable request.

## Code availablility

Not applicable.

