## Supplemental Figures and Tables, and will be used for the link to the file on the preprint site. for "Unveiling differential responses to UVB (305 nm) and UVC (275 nm) in cacao-infecting *Colletotrichum gloeosporioides* and *Pestalotiopsis* sp"

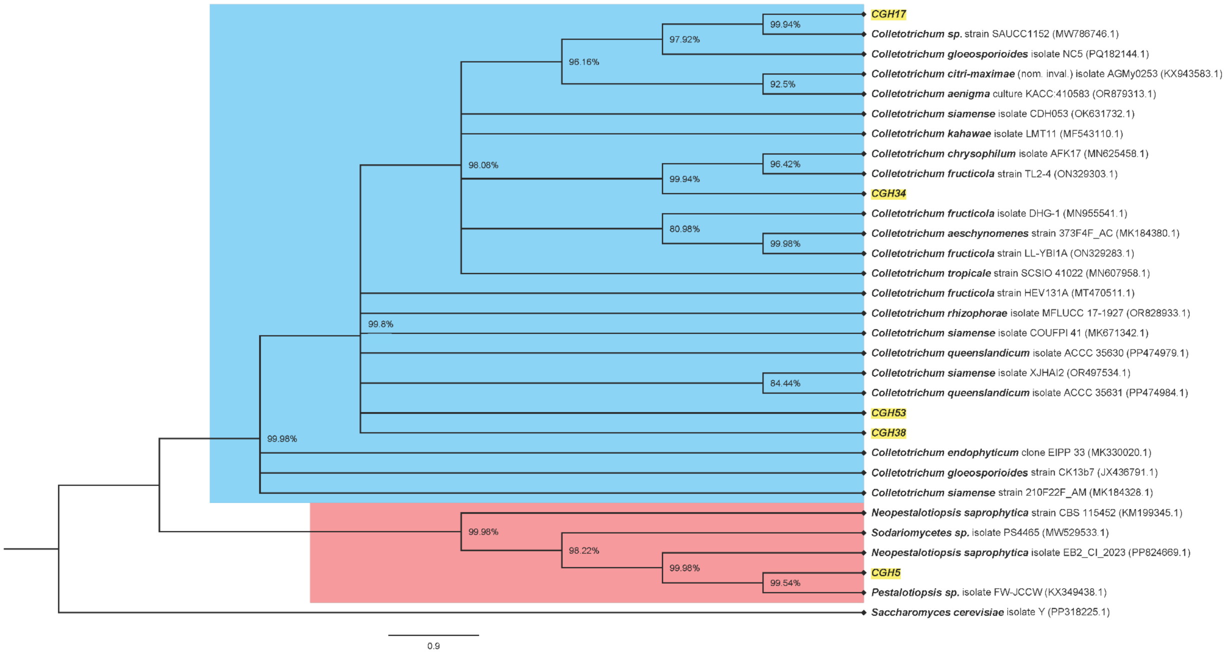

**Fig. S1: Phylogenetic relationships of cacao-associated fungal isolates based on ITS sequence analysis.**

Neighbor-Joining tree constructed using de novo sequences of ITS1 and ITS2 sequences of fungal isolates from cacao and related species obtained from NCBI GenBank. The analysis reveals distinct clades within the *Colletotrichum* genus and places CGH5 within the *Pestalotiopsis* clade. Bootstrap support values (1,000 replicates) are shown at the nodes. The scale bar represents the number of substitutions per site. Supporting values of less than 80% were not included in the tree.

| **Training set** | | **Validation set** | |
|---|---|---|---|
| Method | Misclassification rate | Method | Misclassification rate |
| Bootstrap Forest | 0.1696 | Neural Boosted | 0.2677 |
| Neural Boosted | 0.2651 | Bootstrap Forest | 0.284 |
| Decision Tree | 0.287 | Decision Tree | 0.3022 |
| Support Vector Machines | 0.308 | Nominal Logistic | 0.3205 |
| Nominal Logistic | 0.347 | Generalized Regression Lasso | 0.3631 |
| Generalized Regression Lasso | 0.347 | Support Vector Machines | 0.3773 |
| K Nearest Neighbors | 0.3294 | K Nearest Neighbors | 0.2982 |
| Naive Bayes | 0.5492 | Naive Bayes | 0.5538 |

**Table S1: Performance of machine learning models in classifying fungal isolates based on UV treatment and colony morphology.**

This table compares the accuracy of different machine learning models in predicting five fungal isolates based on UV treatment and colony morphology. Misclassification values indicate the proportion of incorrectly classified isolates. Bootstrap Forest and Neural Boosted achieved over 70% accuracy for training and validation sets.

| Training | Classification accuracy | | | | |
|---|---|---|---|---|---|
| Isolate | CGH5 | CGH17 | CGH34 | CGH38 | CGH53 |
| CGH5 | 0.98 | 0 | 0.002 | 0.017 | 0 |
| CGH17 | 0 | 0.805 | 0 | 0.195 | 0 |
| CGH34 | 0.002 | 0.129 | 0.629 | 0.1 | 0.139 |
| CGH38 | 0.005 | 0.24 | 0.002 | 0.753 | 0 |
| CGH53 | 0 | 0.247 | 0.121 | 0.121 | 0.511 |
| Validation | Classification accuracy | | | | |
| Isolate | CGH5 | CGH17 | CGH34 | CGH38 | CGH53 |
| CGH5 | 0.97 | 0 | 0 | 0.03 | 0 |
| CGH17 | 0 | 0.825 | 0.01 | 0.165 | 0 |
| CGH34 | 0 | 0.176 | 0.549 | 0.099 | 0.176 |
| CGH38 | 0 | 0.25 | 0.019 | 0.731 | 0 |
| CGH53 | 0 | 0.219 | 0.135 | 0.083 | 0.563 |

**Table S2: Performance of Neural Boost (NTanH(3) NBoost(20)) in fungal isolate classification.**

This table presents a detailed comparison of the Neural Boost model's performance in classifying fungal isolates. High accuracy was achieved for CGH5, CGH17, and CGH38 in both training and validation sets, while CGH34 and CGH53 showed comparatively lower accuracy.

| Traits | Main effect | Total effect | Importance |
|---|---|---|---|
| UV treatment | 0.286 | 0.705 | 7 |
| Area size | 0.117 | 0.436 | 4 |
| Length | 0.055 | 0.259 | 2 |
| Width | 0.032 | 0.146 | 1 |
| LWR | 0.025 | 0.129 | 1 |
| Perimeter | 0.02 | 0.116 | 1 |
| Survival rate | 0.02 | 0.106 | 1 |
| Circularity | 0.017 | 0.07 | 0 |
| IS & CG | 0.013 | 0.049 | 0 |

**Table S3: Feature importance in the Neural Boost model for fungal isolate classification.** This table presents the main effect, total effect, and importance of different features in the Neural Boost model used to classify fungal isolates. Main effect represents the direct contribution of a feature, while total effect includes both direct and indirect contributions. UV condition was the most important feature, followed by area size, length, and width. IS & CG, circularity, and survival rate had lower importance in the classification model.

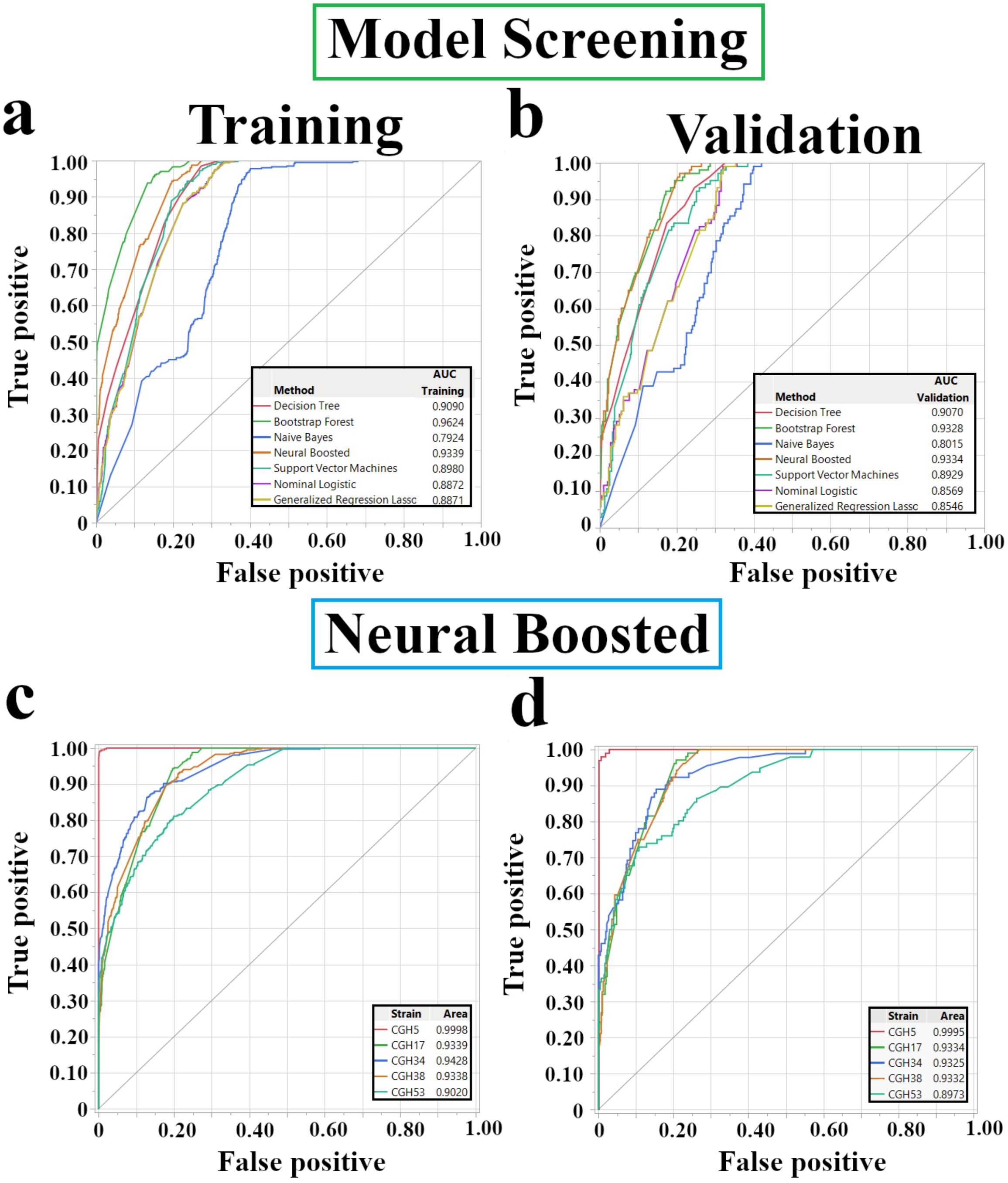

**Fig. S2: Performance evaluation of machine learning models for fungal isolate classification.**

**a-b,** Receiver operating characteristic (ROC) curves for eight machine learning models in classifying fungal isolates based on UV treatment and colony morphology. Higher area under the curve (AUC) values indicate better classification performance. **a,** Training set. **b,** Validation set. **c-d,** ROC curves for the Neural Boosted model, illustrating classification performance for individual fungal isolates. **c,** Training set. **d,** Validation set.

| **Training set** | | **Validation set** | |
|---|---|---|---|
| Method | R-square | Method | R-square |
| Decision Tree | 0.8456 | Boosted Tree | 0.8443 |
| Boosted Tree | 0.8455 | Neural Boosted | 0.8443 |
| Neural Boosted | 0.8448 | Decision Tree | 0.8441 |
| Support Vector Machines | 0.8287 | Support Vector Machines | 0.8206 |
| Fit Least Squares | 0.7910 | Fit Stepwise | 0.7945 |
| Generalized Regression Lasso | 0.7910 | Fit Least Squares | 0.7945 |
| Fit Stepwise | 0.7910 | Generalized Regression Lasso | 0.7945 |
| Bootstrap Forest | 0.7806 | Bootstrap Forest | 0.7939 |
| K Nearest Neighbors | 0.7773 | K Nearest Neighbors | 0.7855 |
| **Boosted tree for area size- Importance of contributions** | | | |
| Trait | Number of splits | Sum of squares | Portion |
| UV treatment | 513 | 144186124 | 0.6519 |
| Isolate | 255 | 76977234 | 0.3481 |
| **Neural Boosted- Feature importance** | | | |
| Trait | Main effect | Total effect | Importance score |
| Isolate | 0.581 | 0.64 | 7 |
| UV treatment | 0.36 | 0.419 | 4 |

**Table S4: Predictive performance of machine learning models for colony area size based on UV treatment and fungal isolate (n = 3,600).**

This table compares the accuracy of nine machine learning models in predicting colony area size using only UV treatment and fungal isolate information (excluding direct size and shape-related traits). Boosted Tree, Neural Boosted, and Decision Tree achieved high accuracy (R-squared > 0.84) in both training and validation sets. The relative importance of UV treatment and fungal isolates varied between models. For Boosted Tree, Importance is measured by the number of splits (representing how often a feature was used to divide the data) and the sum of squares (reflecting the improvement in model accuracy due to a feature). For Neural Boosted, the main effect is the direct contribution of a feature to the prediction, while the total effect includes both the direct and indirect contributions. The importance score is a relative measure of the overall contribution of a feature to the model's predictive accuracy.

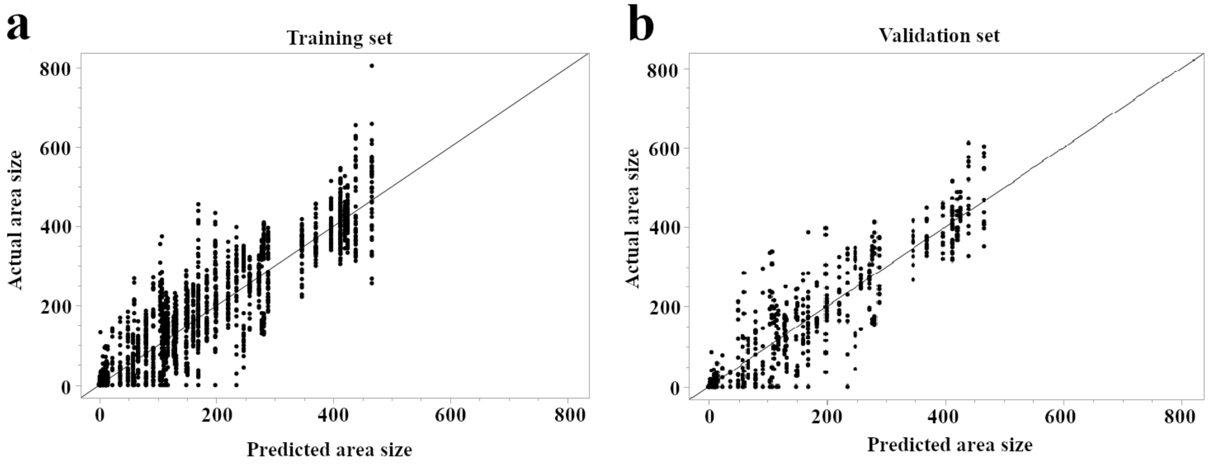

**Fig. S3: Accurate prediction of colony area size using a Neural Boosted model.**

**a-b,** Scatter plots comparing actual versus predicted colony area size values using a Neural Boosted model (n = 3,600). The diagonal line represents perfect prediction (1.0). **a,** Training set (R-squared = 0.8455). **b,** Validation set (R-squared = 0.8443).

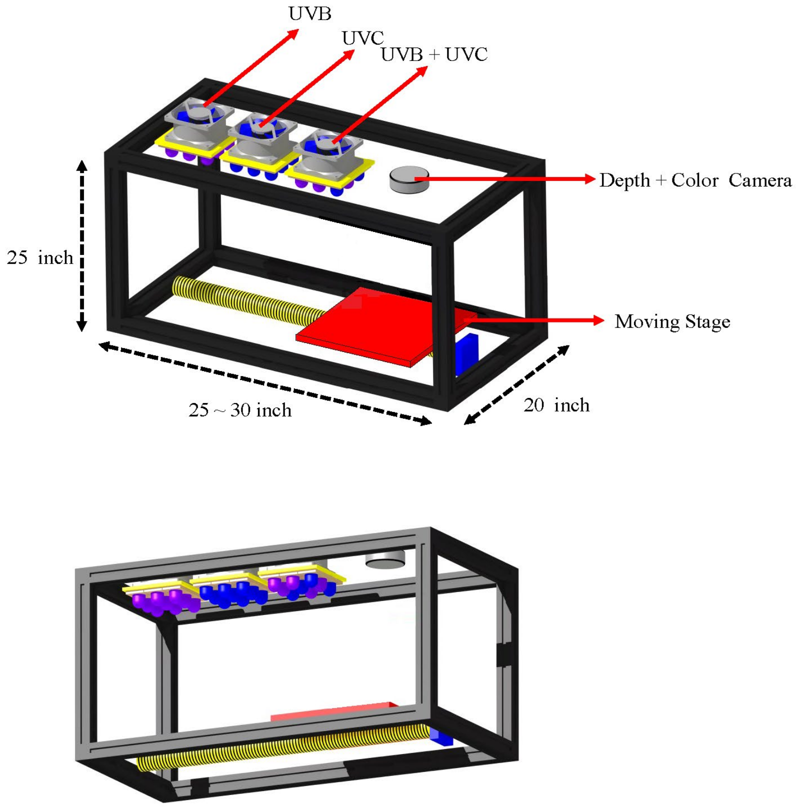

**Fig. S4: Illustration of the second-generation UV treatment system.**

A diagram illustrating the design and key components of the system used for controlled UV exposure of fungal samples. The system features an enclosed frame housing top-mounted LED modules providing UVB, UVC, or combined UVB+UVC radiation. A programmable 'Moving Stage' transports samples through the exposure zone below the LEDs. A Depth + Color camera is mounted alongside the LEDs to capture structural data during treatment. Approximate dimensions are indicated in inches.

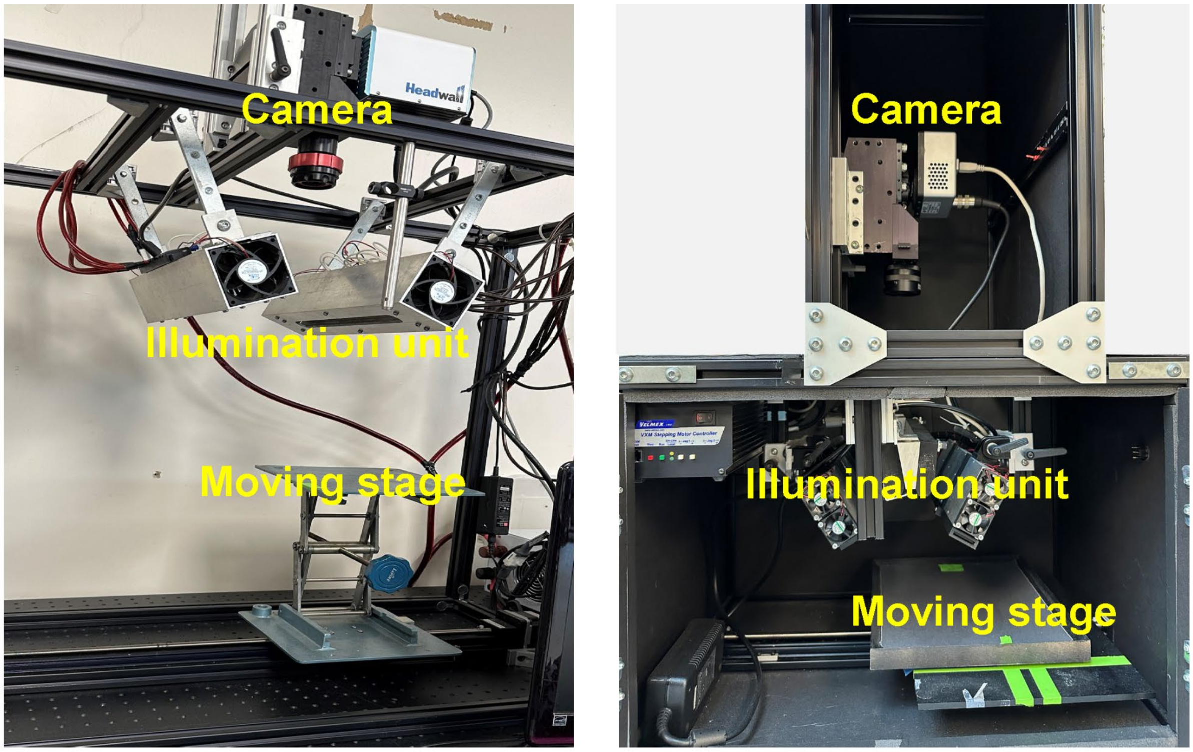

**Fig. S5: Schematic diagrams of the hyperspectral line-scan imaging systems: SWIR system (left) and VIS system (right).**
Each system comprises a spectrograph-coupled camera, a motorized linear stage for sample movement, and customized illumination units optimized for the respective spectral range.
